# LRRC8A/D-containing anion channels mediate swelling-activated taurine release in the mouse brain

**DOI:** 10.64898/2026.10.19.760193

**Authors:** Madison L. Chandler, Julia W. Nalwalk, Preeti Dohare, Ariella D. Sprague, Alexandra L. Schober, Alexander A. Mongin

## Abstract

Volume-regulated anion channels (VRACs) are swelling-activated Cl^−^ channels formed by heteromers of LRRC8 proteins (LRRC8A-E). VRACs mediate regulatory volume decrease by releasing Cl^−^ and organic osmolytes, including taurine. In the brain, taurine functions as both an osmolyte and an inhibitory neuromodulator, acting as an agonist at glycine and GABA_A_ receptors. Here, we used RNAi and genetically modified mice to (i) determine the contribution of LRRC8 subunits to swelling-activated taurine release, and (ii) test whether brain VRACs contribute to systemic osmoregulation. In astrocyte cultures lacking the essential VRAC subunit LRRC8A, swelling-activated [^3^H]taurine release was reduced by ∼90%. Similarly, *in vivo* brain microdialysis revealed a ≥90% reduction in swelling-activated taurine release in brain-wide LRRC8A knockout mice (bLRRC8A KO). RNAi experiments further confirmed LRRC8D as a key determinant of VRAC-mediated taurine release, with *Lrrc8d* knockdown reducing swelling activated [^3^H]taurine efflux by >50%. RNA-seq revealed enrichment of *Lrrc8d* in the hypothalamus, where astrocytic taurine release is proposed to inhibit magnocellular neurons and suppress arginine vasopressin secretion during hypoosmotic conditions. We therefore tested whether disrupting brain VRACs impairs recovery from systemic hypoosmotic stress. bLRRC8A KO mice and mice with gene trap-mediated LRRC8D depletion were challenged with an intraperitoneal injection of sterile water (7.5% body weight, under anesthesia), and serum osmolality was monitored over time. Unexpectedly, neither model showed impaired recovery from hypoosmotic challenge. Overall, this study identifies LRRC8A/D-containing VRACs as the primary conduit for swelling-activated taurine release in the brain, yet indicates that this pathway is not essential for maintaining systemic fluid-electrolyte homeostasis.

## INTRODUCTION

Volume-regulated anion channels (VRACs) are ubiquitously expressed, swelling-activated chloride (Cl^−^) channels that are permeable to a variety of inorganic anions and small organic molecules (1–5). During osmotic cell swelling, VRAC activation promotes regulatory volume decrease (RVD) through the coordinated loss of intracellular ions and organic osmolytes, reducing intracellular osmolarity and driving the efflux of osmotically obligated water (5–7). In addition to their established role in RVD, VRACs provide an important pathway for the release of various signaling molecules, including the neuroactive compounds glutamate, GABA, taurine, and ATP [see (8–15) and reviews (16–19). VRACs are formed as heteromeric assemblies of proteins belonging to the leucine-rich repeat-containing family 8 (LRRC8A-E), with LRRC8A serving as an essential subunit (20, 21). Structural studies have shown that LRRC8 proteins assemble into hexameric channels with one central ion-conducting pore (22–28). LRRC8A must assemble with one or more additional LRRC8 isoforms to form a functional channel, and the resulting subunit composition determines key VRAC properties, including substrate selectivity (13, 21, 29–32). For example, LRRC8A/LRRC8D heteromers exhibit high permeability to neutral organic osmolytes, including myo-inositol, GABA, and taurine (13, 31, 33).

Taurine (2-aminoethanesulfonic acid) is an atypical sulfur-containing amino acid and one of the most abundant organic osmolytes in mammalian tissues, including the brain (34, 35). In addition to its established roles in cell volume regulation and cytoprotection, taurine serves as an endogenous inhibitory neuromodulator (36). Extracellular taurine regulates neuronal excitability by acting as a partial agonist at glycine and GABA_A_ receptors (37–39). In the brain, taurine is enriched in neurons, astrocytes, and endothelial cells, but its cell-specific distribution varies by region (40). Astrocytes maintain high intracellular taurine concentrations through both uptake via the sodium-dependent taurine transporter (TauT, encoded by *Slc6a6*) and synthesis from cysteine through the intermediate hypotaurine (41, 42). Neurons have lower taurine synthesis capacity and may, therefore, depend on astrocytic supplies of hypotaurine and taurine (43). Astrocytes represent an important reservoir of taurine, and can release it in response to various stimuli, including cell swelling (8, 12, 44). Thus, both swelling- and agonist-induced VRAC activation can cause taurine release, providing a mechanism by which VRACs may modulate neuronal excitability.

VRAC-mediated taurine release is particularly relevant in hypothalamic circuits which regulate systemic fluid and electrolyte homeostasis (45, 46). In the hypothalamus, changes in plasma osmolarity are detected by osmosensitive neurons in several periventricular areas, including the median preoptic nucleus, the organum vaculosum lamina terminalis, and the subfornical organ, which project to the paraventricular (PVN) and supraoptic (SON) nuclei (46, 47). Magnocellular neurons in the PVN and SON synthesize and secrete the antidiuretic hormones arginine vasopressin (AVP) and oxytocin, which are released from the posterior pituitary (46, 47). Hyperosmotic conditions increase magnocellular neuronal activity and AVP release to promote water retention, whereas hypoosmotic conditions suppress AVP secretion and facilitate water excretion. Within these circuits, astrocyte-derived taurine has been proposed to modulate neuronal activity (45, 48, 49). Specifically, reductions in extracellular osmolarity are thought to activate astrocytic VRACs, promoting taurine release that inhibits magnocellular neurons through glycine receptor activation and suppresses pituitary AVP secretion (37, 48, 49). Thus, taurine signaling is thought to represent a critical brake on AVP and oxytocin release.

In the present study, we used both an RNAi approach and genetically modified primary mouse astrocytes to determine the LRRC8 subunit requirements for endogenous taurine-permeable VRACs. We further assessed the contribution of taurine-permeable VRACs to systemic osmoregulation by measuring serum osmolality following acute hyperosmotic and hypoosmotic challenges in genetically modified mice with targeted deletion of LRRC8A or LRRC8D.

## MATERIALS AND METHODS

### Ethical approval

All animal procedures used in this study were approved by the Institutional Animal Care and Use Committee of Albany Medical College (ACUP 24-03002), and were conducted in accordance with the Guide for the Care and Use of Laboratory Animals as adopted by the U.S. National Institutes of Health (https://grants.nih.gov/grants/olaw/guide-for-the-care-and-use-of-laboratory-animals.pdf).

### Animals

This study used three mouse models for *in vitro* and *in vivo* analyses. All animals were housed in a facility with controlled temperature, humidity, and lighting, maintained on a 12 h light/dark cycle, and provided free access to food (Purina Lab Diet 5P76) and water. Experiments included age-matched experimental and control mice of both sexes, with males and females represented as evenly as possible. Experiments are reported in accordance with the ARRIVE guidelines. No animals were lost to attrition during the study period.

Wild type (WT) animals for RNAi-based experiments were of the C57BL/6J background, obtained from Jackson Laboratory (Farmington, CT, USA; strain #000664, RRID: IMSR_JAX:000664) and bred in-house.

Mice with conditional deletion of *Lrrc8a* in the brain (bLRRC8A KO) were produced by breeding *Lrrc8a*^flox/flox^ females (gift of Dr. Rajan Sah, Washington University School of Medicine, St. Louis, MO, USA) with commercially available *Nestin*^Cre/+^ males (B6.Cg-Tg(Nes-cre)1Kln/J; Jackson Lab stock #003771, RRID:IMSR_JAX:003771). The resulting heterozygous *Nestin*^Cre/+^:*Lrrc8a*^flox/+^ males were then crossed with *Lrrc8a*^flox/flox^ females to produce *Nestin*^Cre/+^:*Lrrc8a*^flox/flox^ mice (bLRRC8A KO) and littermate controls. The specificity and efficacy of this genetic strategy were validated in our previous work (32, 50, 51). To visualize Cre activity and assess potential unintended germline recombination, mice were crossed with a strain carrying the Ai9 tdTomato Cre reporter (B6.Cg-Gt(ROSA)26Sor^tm9(CAG-tdTomato)Hze^/J; Jackson Lab stock #007909, RRID:IMSR_JAX:007909). Due to the lethal phenotype of bLRRC8A KO animals during late adolescence, all *in vivo* studies were conducted between 6-8 weeks of age (50).

Commercially available, “gene-trapped”, *Lrrc8d* knockout mice were purchased from Taconic (Model TF2126, cryopreserved; Germantown, NY). Heterozygous *Lrrc8d*^-/+^ mice, on a 129/SvEv-C57BL/6 background as provided, were bred as pairs, and further inbred into a C57BL/6J background. This breeding strategy results in three genotypes, confirmed by PCR analysis as: *Lrrc8d*^+/+^ (wild type, WT), *Lrrc8d*^-/+^ (heterozygous deletion, Het), and *Lrrc8d*^-/-^ (constitutive knockout, KO). Insertion of the trapped construct into mouse chromosome 5 was validated according to the Taconic genotyping protocol.

### Brain tissue collection and processing

For animals from the bLRRC8A KO line, mice of both sexes were euthanized at 6-8 weeks of age due to the lethal phenotype associated with bLRRC8A deletion (50). Animals of both sexes from the gene-trapped LRRC8D KO line were euthanized at 8-10 weeks of age. Mice were euthanized with a lethal intraperitoneal (i.p.) injection of sodium pentobarbital (100 mg/kg) then transcardially perfused with 20 mL room temperature phosphate-buffered saline (PBS), followed by 20 mL ice-cold PBS. Brains were immediately removed, and the olfactory bulbs and cerebellum were dissected away. The remaining brain tissue was divided into two hemispheres and immediately snap-frozen in liquid nitrogen. Samples were stored at −80^◦^C until processed.

Tissue for Western blot analysis was homogenized in lysis buffer containing 150 mM NaCl, 50 mM Tris-HCl (pH 8.0), 0.1% Triton X-100, and 5% protease inhibitor cocktail (Thermo Fisher Scientific, Waltham, Massachusetts, USA, cat. #78430) using a mechanical homogenizer (PRO200, PRO Scientific, Oxford, CT, USA). Homogenates were clarified by centrifugation (10,000 g for 5 min at 4°C) and the supernatants were collected. Protein concentration was determined using the Pierce Bicinchoninic acid protein assay kit (BCA, Thermo Fisher, cat. #23225). The remaining lysate was diluted with 2× reducing Laemmeli buffer (Bio-Rad Laboratories, Hercules, CA, USA, cat. #1610737) and stored at −80^◦^C until processed.

For high-performance liquid chromatography (HPLC), one hemisphere of snap-frozen brain tissue was homogenized in 0.4 M perchloric acid. Homogenates were clarified by centrifugation (10,000 g for 10 min), and the resulting supernatants were collected for determination of tissue amino acid concentrations by HPLC. The pellets were saved to determine total protein content and normalize amino acid concentrations.

### Primary cortical astrocyte cultures and sample preparation

Primary cultures of mouse astrocytes were prepared from the brain cortices of newborn (P0-P2) animals of both sexes according to an institutionally approved protocol. Neonates were anesthetized by cooling, rapidly decapitated, and the brains were transferred to ice-cold sterile Dulbecco’s phosphate-buffered saline without calcium and magnesium (Thermo Fisher, cat. #14190144). The cerebral cortices were cleared of meninges and isolated. Tissue was minced using a sterile surgical blade, digested for 5 min at room temperature (22°C) using 500 µL of TrypLE Express recombinant protease (Thermo Fisher, cat. #12605), and additionally mechanically triturated by pipetting. Enzymatic digestion was terminated by adding Earl’s minimal essential medium (MEM, Thermo Fisher, cat. #11095), supplemented with 10% heat inactivated horse serum (HIHS, Thermo Fisher, cat. #26050), and penicillin/streptomycin (P/S, 10,000 U/mL and 10,000 µg/mL, respectively, ThermoFisher, cat. #15140). Next, the cell suspension was filtered through a 70 µm nylon mesh cell strainer (Corning, Corning, NY, USA, cat. # 431751). Cells from the cortices of half brains (WT) or full brains (bLRRC8A KO and “gene-trapped” LRRC8D) were plated onto T75 flasks to obtain enriched astrocytic populations. T75 flasks were kept at 37°C in a humidified atmosphere containing 5% CO_2_/balance air. Cells were re-fed biweekly with fresh MEM+HIHS+P/S medium. Primary cultures reached confluency after ∼2 weeks and were maintained for up to six weeks until re-plating for analysis.

Western blot samples were prepared from confluent 60 mm Petri dishes, washed of cell culture media with 1x DPBS, and lysed with 2% sodium dodecyl sulfate (SDS) plus 8 mM EDTA. The 60 mm dishes were pretreated with Poly-D-Lysine (PDL) to facilitate cell attachment.

Protein concentration was determined using the Pierce Bicinchoninic acid protein assay kit (Thermo Fisher, cat. #23225). The remaining lysate was diluted with 4× reducing Laemmeli buffer (Bio-Rad, cat # 1610747) and stored at −80^◦^ until analysis.

### Western blot analyses

Protein lysates from primary astrocyte cultures and forebrains were prepared as described above. Samples in Laemmeli buffer were boiled for 5 min, and 10 or 20 µg of total protein per sample was resolved on 10% Mini-PROTEAN TGX gels (Bio-Rad, cat #4561033) using standard SDS-PAGE procedures. Separated proteins were electro-transferred onto a polyvinylidene difluoride membrane (PVDF, Bio-Rad, cat. #1620177) using a Trans-Blot Turbo transfer system (Bio-Rad, RRID:SCR_023156). Membranes were blocked for 5 min in Tris-buffered saline containing 0.1% Tween-20 (TBS-T) and 5% milk. They were further incubated overnight in the same blocking buffer with primary antibody at 4°C. Blots were washed with TBS-T and probed with the species-matching secondary horseradish peroxidase-conjugated antibody in TBS-T containing 5% milk for 2 h at room temperature. After a final wash, immunoreactivity was measured using ECL reagent (GE Healthcare, Chicago, Illinois, USA, cat. #RPN2232) and visualized with a ChemiDoc Imager (Bio-Rad, RRID:SCR_019684). For loading controls, membranes were washed overnight in TBS-T and re-probed with the HRP-conjugated anti-β-actin antibody for 20 min and β-actin immunoreactivity was visualized as described above. Band intensities were quantified using ImageJ software (NIH, RRID:SCR_003070) and were normalized to β-actin immunoluminescence from the same blot. Detailed information on all antibodies and their dilutions is provided in Table 1.

**Table 1.** Primary and secondary antibodies used for western blot and immunohistochemical analysis.

| Protein target | Species/type | Source | Catalog # | RRID | Dilution |
| --- | --- | --- | --- | --- | --- |
| β-actin, HRP-conjugated | Mouse monoclonal | Millipore-Sigma | A3854 | AB_262011 | 1:50,000 |
| LRRC8A | Mouse monoclonal | Santa Cruz Biotech. | sc-517113 | AB_2928142 | 1:1,000 |
| LRRC8C | rabbit polyclonal | Proteintech | 21601-1-AP | AB_10733648 | 1:2,000 |
| LRRC8D | rabbit polyclonal | Proteintech | 11537-1-AP | AB_2137767 | 1:2,000 |
| c-Fos | Rabbit monoclonal | Cell Signaling Technology | 2250 | AB_2247211 | 1:1,000 |
| Secondary (WB): anti-mouse IgG, HRP-conjugated | sheep | GE Healthcare | NA931 | AB_772210 | 1:10,000 |
| Secondary (WB): anti-rabbit IgG, HRP-conjugated | donkey | GE Healthcare | NA934 | AB_772206 | 1:10,000 |
| Secondary (IHC): goat anti-rabbit IgG, Alexa-Fluor 594 | goat | Thermo Fisher Scientific | A32740 | AB_2762824 | 1:400 |

In several instances the same membranes were reprobed with primary antibodies recognizing different LRRC8 proteins when raised in different species (e.g., mouse anti-LRRC8A and rabbit anti-LRRC8C). This is clearly indicated in figure legends. The complete dataset of analyzed western blots, including matching β-actin loading controls, is included in Supplemental Materials.

### VRAC activity assays

VRAC activity was assessed by measuring the swelling-activated release of either [^3^H]taurine or D-[^14^C]aspartate (a non-metabolizable analog of glutamate), as extensively validated in our previous studies (31, 52, 53). Astrocytes were cultured on PDL-treated 18 × 18 mm glass coverslips and preloaded overnight in standard serum-containing medium supplemented with 2 µCi/mL [^3^H]taurine (Perkin Elmer, Waltham, MA, USA) or D-[^14^C]aspartate (American Radiolabeled Chemicals, Inc., St. Louis, MO, USA).

To characterize the kinetics of VRAC activation, radiotracer efflux was measured using a custom-fabricated Lucite superfusion chamber with a recessed base to accommodate an 18 × 18 mm coverslip and a Teflon screw lid that maintained a ∼200 µm gap above the cell monolayer. Coverslips were briefly rinsed with isoosmotic basal medium containing (in mM): 135 NaCl, 3.8 KCl, 1.2 MgSO_4_, 1.3 CaCl_2_, 1.2 KH_2_PO_4_, 10 D-glucose, and 10 HEPES (pH 7.4; osmolality 290 ± 3 mosmol/kg H_2_O). Coverslips were then transferred to the superfusion chamber, and continuously perfused at a flow rate of ∼1.4 mL/min with either isoosmotic or hypoosmotic medium in which NaCl was reduced by 50 mM (∼200 mosmol/kgH_2_O; ∼30% reduction) or 80 mM (∼150 mosmol/kgH_2_O; ∼50% reduction) to induce cell swelling and activate VRACs. One-minute perfusate fractions were collected using an automated CF-1 fraction collector (SpectraLab Scientific, Markham, ON, Canada). At the end of each experiment, cells were lysed in 2% SDS plus 8 mM EDTA. Scintillation liquid (Ecoscint, Atlanta Biologicals, Atlanta, GA, USA, cat. #LS-273) was added to the samples and radiotracer content in individual samples was determined using a TriCarb 4910 TR scintillation counter (Perkin Elmer, Shelton, Connecticut). Data were presented as fractional release values, calculated at each time point as the ratio of released radiotracer to total radiotracer content (cumulative release plus remaining intracellular tracer), using a custom Excel-based analysis program.

### *In vivo* brain microdialysis

For microdialysis experiments, 6-8-week-old male and female mice of either the *Lrrc8a*^fl/+^ (control) or bLRRC8A KO genotype were used. Anesthesia was induced with 5% isoflurane in 30% O_2_/balance N_2_ and maintained throughout surgical procedures with 1.25 to 1.5% isoflurane under constant observation. To prevent excessive fluid secretion in the respiratory system and gastrointestinal tract, 0.04 mg/kg of glycopyrrolate was given via subcutaneous injection. Animals were kept hydrated with injections of 0.2 cc saline i.p. three times while under anesthesia. Body temperature was maintained at 36.5°C using a heating pad and monitored with a probe placed rectally (RightTemp Jr., Kent Scientific, Torrington, CT, USA). Mice were placed in a stereotaxic frame (David Kopf Instruments, Tujunga, CA) and the probe guide introduced through a small burr hole in the skill. Dental cement (3M Durelon Carboxylate Luting Cement; Solventum, Eagan, MN, USA) and 3 small screws (000-120; JI Morris, Southbridge, MA, USA) were used to anchor the assembly to the skull. A single microdialysis probe (CX-I series, 0.2232 mm membrane, 50,000 kD molecular weight cutoff, 2 mm membrane length; Eicom Corporation, San Diego, CA) was lowered into the striatum through the guide (target placement for tip of probe 0.5 mm AP, 1.8 mm ML, and -5 mm DV relative to bregma; Paxinos and Franklin, 2001).

Artificial cerebrospinal fluid (aCSF) was perfused at a flow rate of 2 µl/min throughout the experiment. The composition of isoosmotic aCSF was as follows (in mM): 120 NaCl, 2.7 KCl, 1 MgSO4, 1.2 CaCl2, 25 NaHCO3, 0.05 ascorbic acid (pH=7.3). Following a 1 h baseline stabilization period, microdialysis samples were collected every 20 min for 2 h using a CMA 470 refrigerated fraction collector (CMA Microdialysis, Holliston, MA). The solution flowing through the probe was then switched to hypoosmotic aCSF (H-aCSF, 2 µl/min, 25 mM NaCl; all other components remained the same as above) and fractions collected every 5 minutes for 1 h. Finally, isoosmotic aCSF was again perfused through the probe for 1 hr and fractions collected every 20 min. All samples were kept at 4^◦^C prior to same day, off-line, determination of amino acid levels using HPLC.

### High-performance liquid chromatography (HPLC)

Amino acid levels were measured by reverse-phase HPLC using an Agilent 1200 HPLC setup and Eclipse XDB-C18 column (both from Agilent Technologies, Santa Clara, CA, USA) as described in (50, 51). Briefly, precolumn derivatization was performed with a freshly prepared mix of o-phthalaldehyde and 2-mercaptoethanol in 0.4 M sodium tetraborate buffer (pH = 9.5). The amino acid derivatives were eluted with a mobile phase solvent containing 30 mM NaH_2_PO_4_, 30 mM sodium acetate, 2% methanol, 1% tetrahydrofuran, 0.05% sodium azide, in combination with varying concentrations of HPLC grade methanol (10-30%). Signal was measured using a programmable 1200 series fluorescence detector (Agilent, Lexington, MA, USA). Amino acid standards were processed in the same fashion and used to identify amino acid peaks and calculate concentrations of individual amino acids in the samples.

### Computational analysis of brain RNA-seq data from the human protein atlas

Publicly available RNA-seq data were obtained from the Human Protein Atlas (HPA) and analyzed using R (v4.3.1; RRID; R Foundation for Statistical Computing, Vienna, Austria) and RStudio (v2023.12.1+402; RRID; Posit Software, Boston, MA, USA). Mouse brain expression data were obtained from the HPA version 22.0 mouse brain sample dataset (Ensembl version 103.38; rna_mouse_brain_sample_hpa.tsv.zip), which contains *Lrrc8a-e* transcript expression data from 13 brain regions collected from two male and two female C57BL/6N mice (2 months of age). Data were downloaded on April 29, 2026. Human brain expression data were obtained from the HPA version 25.1 brain region dataset (Ensembl version 109; rna_brain_region_hpa.tsv.zip), which contains transcript expression levels summarized across 13 major brain regions. Data were downloaded on September 16, 2026. The HPA brain RNA-seq datasets were generated as part of the mammalian brain mapping project described by Sjöstedt et al. (54). The Human Protein Atlas datasets were accessed at the version-specific HPA website (https://www.proteinatlas.org/about/download). Additional datasets of publicly available human RNA-seq data were obtained from the Genotype Tissue Expression project (GTEx, https://gtexportal.org) on September 29, 2026.

### siRNA-mediated knockdown

Knockdown of specific *Lrrc8* transcripts was performed using siRNA constructs from Qiagen (Valencia, CA, USA), as described in our previous work (32). Briefly, primary WT astrocytes were grown to 70-80% confluency on PDL-treated 60 mm dishes and transfected with siRNA using Lipofectamine RNAiMAX (Thermo Fisher, cat. # 13778-075). siRNA/Lipofectamine complexes were prepared in Opti-MEM as per the manufacturer’s protocol, further diluted with Opti-MEM to a final siRNA concentration of 20-50 nM (optimized for individual siRNAs) and added to the cells. After 4 h, cultures were supplemented with MEM containing 10% HIHS and penicillin plus streptomycin (MEM+HIHS+P/S). At 24 h, the transfection medium was completely replaced with fresh MEM+HIHS+P/S. As a control, we used the scrambled AllStars negative control siRNA (Qiagen, Germantown, Maryland, USA, cat. # SI03650318). Changes in mRNA expression were assessed by qRT-PCR 48 h after transfection (32), whereas protein expression and functional assays were evaluated 96 h after transfection. Details of the siRNA constructs are listed in Table 2.

**Table 2.**
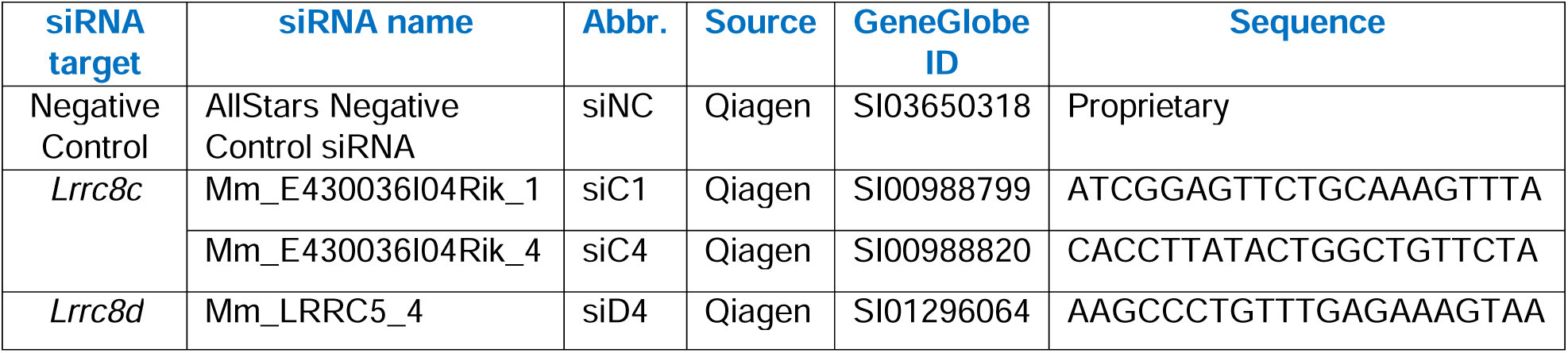
siRNA constructs used in RNAi experiments.

### *In Vivo* osmotic challenge experiments

To explore potential changes of systemic osmoregulation in LRRC8D KO mice, we exposed animals from all 3 genotypes (WT, Het, and KO) to either hyper- or hypoosmotic challenge and measured serum osmolality at several time points post-challenge. Mice of both sexes were used for all time points and treatments. Due to availability in the colony, animals were tested between 8 to 18 wks of age, with most animals tested at approximately 9 wks of age (9.3 ±0.2 wks, n=133). Animals, in home cages, were brought up to the test area and given 30 min to habituate prior to testing. During the osmotic challenge, animals were anesthetized with 5% isoflurane to limit stress responses and hormone release during handling.

The first cohort of ‘no injection’ controls were weighed and immediately sacrificed while under deep anesthesia; their trunk blood was collected into a serum separation tube (SST, cat. #367983, BD Biosciences, Franklin Lakes, NJ, USA). The second cohort of anesthetized mice were subjected to hyperosmotic challenge: they were weighed and injected i.p. with 2N NaCl (20 µl/g body weight) while under anesthesia. Mice were then returned to the home cage, and water access was restricted for the first 2 h post-injection, but then fully restored. At 2, 6 and 24 h post injection, mice were re-anesthetized, rapidly decapitated, and their trunk blood was collected. The third cohort of anesthetized mice were subjected to a hypoosmotic challenge and received an i.p. injection of sterile water (7.5% body weight). These animals had free access to water throughout the experiment. They were anesthetized and euthanized at 30 min, 1 h, or 2 h after water injection. Following collection of trunk blood, all SST samples rested at room temperature for 2 h and were then spun at 2500 g for 15 min. Serum was collected, and osmolality measured using a freezing point osmometer (µOsmette, Precision Systems, Natick, MA).

The hypoosmotic challenge experiments were repeated in bLRRC8A KO animals, which exhibit complete loss of VRAC activity in the brain. The bLRRC8A KO group had a smaller sample size due to the lethal phenotype of this transgenic model, which limited the number of animals surviving to the testing age (50). Because of the rapid recovery of systemic osmolarity observed in the LRRC8D-deficient cohort following water injection, we limited the bLRRC8A KO post-injection observation period to 1 h. In bLRRC8A mice, we additionally quantified systemic taurine content in blood serum using HPLC. For HPLC analysis, a fixed volume of 50 µL serum was used for each sample. Proteins were precipitated by adding 0.8 N HClO_4_ at a 1:1 ratio with the serum sample. Samples were then centrifuged at 10,000 g for 6 min, and the resulting supernatant was collected. Prior to HPLC processing, the sample pH was adjusted by addition of 10 N NaOH, and each sample was subsequently diluted 15-fold with Milli-Q water before analysis.

### Immunohistochemistry

Mice of both sexes, aged 9-20 weeks, were used for immunohistochemical analysis based on availability within the colony. Animals were euthanized with sodium pentobarbital (100 mg/kg, i.p.) then transcardially perfused with 20 mL each of room-temperature PBS, ice-cold PBS, and 4% paraformaldehyde (PFA) in PBS. Brains were removed, postfixed in 4% PFA for 24 h, transferred to a 15% sucrose solution for 24 h, followed by a 30% sucrose solution for 24 h for full cryoprotection; all steps were performed at 4°C. Tissue was frozen to -20°C, sectioned at 25 μm (Leica CM3050 cryostat, Wetzlar, Germany; RRID: SCR_016844) and stored in cryoprotectant solution at -20°C (30% ethylene glycol and 20% glycerol in Tris-buffered saline; TBS).

For immunohistochemical staining, free-floating sections were washed with PBS (one 15 min wash followed by four 5 min washes) and blocked for 1 h at room temperature in PBS containing 5% goat serum and 0.2% Triton X-100. Sections were then incubated overnight at 4°C in the blocking solution containing rabbit monoclonal anti-c-Fos primary antibody (see Table 1). The next day, sections were washed with PBS and incubated for 2 h at room temperature with the secondary antibody goat anti-rabbit Alexa-Fluor 594 (see Table 1) and DAPI (0.05 μg/mL; Millipore Sigma, Burlington, MA, USA, cat. # 32670). Following additional PBS washes, sections were mounted onto glass slides, dried overnight at room temperature in the dark, and then cover slipped using ProLong Glass Antifade Mountant (Thermo Fisher, cat. no. P36984).

Fluorescently labeled sections were imaged using a Zeiss Axio Imager M2 microscope (RRID: SCR_024706) at 200× magnification, with image acquisition settings kept consistent across samples. Individual images were stitched using Neurolucida software, version 11.03 (MBF Bioscience; RRID: SCR_001775).

### Statistics

All data are presented as mean values ± SD, with the number of independent experiments (n) indicated. For all *in vivo* experiments, n represents an individual animal. In radiotracer efflux assays, n represents an individual experiment with biological replicates from at least two different cell cultures prepared from different animals. Due to the exploratory nature of this work, power calculations were performed for some, but not all presented experiments. Generally, we planned to be able to detect a difference of 25% between groups with a power of 0.8 and an α of 0.05.

Experimental data were tested for normal distribution. Statistical comparisons between two groups were performed using an unpaired t-test. Comparisons among three or more groups were performed using one-way ANOVA followed by Tukey’s post hoc multiple-comparisons test. Osmotic challenge experiments were analyzed using two-way ANOVA, with main effects of time and genotype. When significant main effects or interactions were detected, Tukey’s post hoc multiple-comparisons test was used to identify differences between groups. Body weights of gene-trapped LRRC8D mice were analyzed using two-way ANOVA, with main effects of sex and genotype.

All statistical analyses and graphical preparations were carried out using GraphPad Prism 10.0 (GraphPad Software, San Diego, CA, USA; RRID:SCR_002798). Full statistical analyses are provided in supplemental information deposited in Open Science Framework. All data have been included in the primary data files for each figure, and no data has been excluded from analysis.

## RESULTS

### LRRC8A-containing VRACs are the main conduit for swelling-activated taurine release in primary astrocytes

To determine the contribution of VRACs to swelling-activated taurine release, we cultured primary astrocytes from mice with a brain-wide knockout of the essential VRAC subunit LRRC8A. Conditional brain recombination was achieved by crossing *Lrrc8a* floxed mice with Nestin^Cre^ animals, resulting in gene deletion of *Lrrc8a* in all cells of neuroectodermal origin, including neurons, astrocytes, and oligodendrocytes (55, 56). This breeding scheme generated two floxed control genotypes (*Lrrc8a*^fl/+^ and *Lrrc8a*^fl/fl^; referred to as fl/+ and fl/fl), a heterozygous deletion (HET), and a homozygous brain-wide knockout (bLRRC8A KO). The bLRRC8A KO model has been extensively characterized in our previous studies and exhibits complete loss of LRRC8A protein in the brain and in primary astrocytes derived from these animals (32, 50, 51). To ensure that the new set of transgenic mice used in the present work retained full LRRC8A loss, we performed additional western blot analysis of forebrain protein lysates. We found a ∼50% reduction in LRRC8A protein in HET animals relative to controls (p<0.0001, Fig. 1A), whereas LRRC8A immunoreactivity was undetectable in bLRRC8A KO brains (p<0.0001; Fig. 1A; full Western blot dataset in Supplemental Fig. S1).

**Figure 1:**
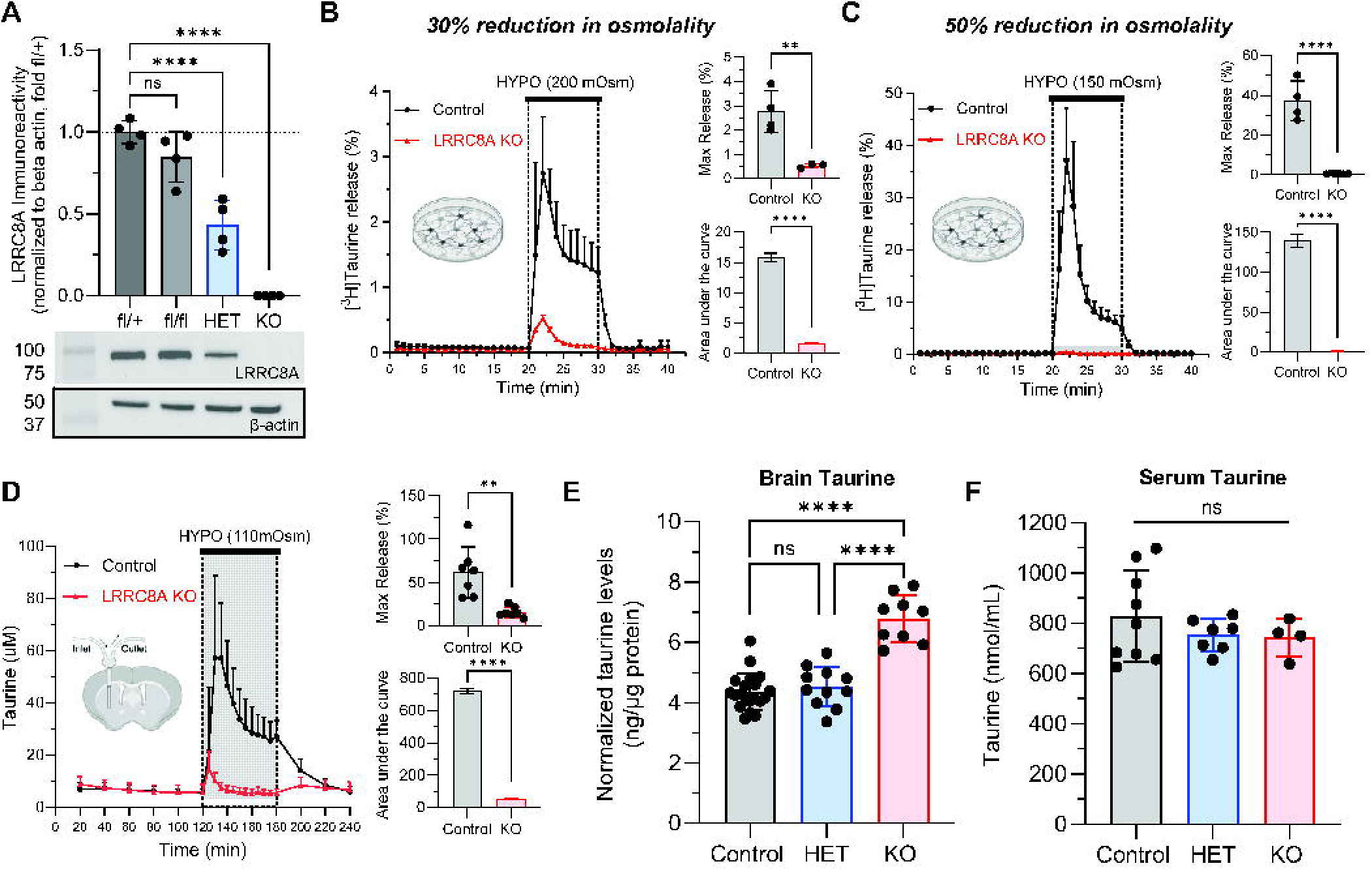
LRRC8A-containing VRACs are the primary pathway for swelling-activated taurine release in astrocytes and in the brain. **A**, Quantification of LRRC8A protein levels in forebrain lysates from the four genotypes resulting from the Nestin^Cre^-driven *Lrrc8a* deletion in the brain: *Lrrc8a*^flox/+^ (fl/+), *Lrrc8a*^flox/flox^ (fl/fl), *Lrrc8a*^flox/+^:*Nestin*^Cre/+^ (HET), and *Lrrc8a*^flox/flox^:*Nestin*^Cre/+^ (KO). Data are the mean values ± SD from six animals/group normalized to an average fl/+ control. ****p<0.0001, one-way ANOVA with Tukey’s multiple comparisons test. Bottom panel shows representative western blot of LRRC8A immunoreactivity and corresponding β-actin signal. **B**, VRAC activity in primary astrocytes assessed as swelling-activated release of [^3^H]taurine in response to a 30% reduction in medium osmolality. Data are the mean values ± SD from 3-4 independent experiments in at least one astrocyte culture per group. *Insets*, Statistical comparison of the maximal (top) and integral (bottom) [^3^H]taurine release values in panel *B*. ****p<0.0001, **p < 0.01, unpaired t-test. **C**, VRAC activity in primary astrocytes assessed as swelling-activated release of [^3^H]taurine in response to a 50% reduction in medium osmolality. Data are the mean values ± SD from 4-6 independent experiments in two different astrocyte cultures per group. *Insets*, Statistical comparison of the maximal (top) and integral (bottom) [^3^H]taurine release values in panel *C*. ****p<0.0001, unpaired t-test. **D**, Microdialysis measurements of extracellular taurine in the mouse striatum to following exposure to hypoosmotic aCSF. Data are the mean values ± SD from seven animals per genotype. *Insets*, Statistical comparison of the maximal (top) and integral (bottom) taurine release values in panel *D*. ****p<0.0001, **p<0.01, unpaired t-test. **E**, Taurine levels measured by HPLC in forebrain lysates from control (fl/+ and fl/fl), HET, and bLRRC8A KO animals, normalized to protein content. Data are the mean values ± SD of 9-20 independent animals per genotype. ****p<0.0001, one-way ANOVA with Tukey’s multiple comparisons test. Adapted with permission from (50). **F**, Serum taurine levels measured by HPLC from trunk blood in the same genotypes. Data are the mean values ± SD of 4-9 independent animals per genotype, one-way ANOVA.

We first assessed VRAC contributions to taurine release *in vitro* using a [^3^H]taurine efflux assay in primary mouse astrocytes. In control cells, a 30% reduction in extracellular osmolality (to 200 mosmol/kgH_2_O) induced a rapid increase in taurine release, which reached a peak rate of 2.7% within 2 min (a 34-fold rise over baseline) (Fig. 1B). This kinetic profile closely matches VRAC activation in both rat and mouse astrocytes measured in our prior studies (31, 32, 52). In contrast, LRRC8A-null astrocytes showed a profound loss of swelling-induced taurine release, with a ≥80% reduction in peak release (p=0.007; Fig. 1B) and a ∼90% decrease in total (integrated) release under hypoosmotic conditions (p<0.0001; Fig. 1B) compared with controls. Interestingly, LRRC8A KO astrocytes also showed a significant reduction in baseline taurine release under isoosmotic conditions: on average 48% lower than that of littermate controls (p=0.012, Supplemental Fig. S2C). A small, transient taurine release persisted during the initial 3 min, perhaps due to activation of VRAC-independent pathway or cell lysis (Fig. 1B, expanded scale in Supplemental Fig. S2A).

To evaluate whether this residual taurine release in LRRC8A-null astrocytes was sensitive to the magnitude of osmotic challenge, we exposed astrocytes to a more severe 50% reduction in extracellular osmolality. In control cells, when medium osmolality was reduced to 150 mosmol/kgH_2_O, VRAC activity was strongly enhanced, with taurine release reaching a maximum release rate of 37.1% within 2 min (Fig. 1C). This represents a 13.7-fold increase relative to the 30% osmotic reduction condition (compare Fig. 1B and Fig. 1C). In contrast, LRRC8A-null astrocytes again showed a near-complete loss of swelling-induced taurine release, with >98% reduction in peak release (p<0.0001; Fig. 1C) and >99% reduction in integral release under hypoosmotic conditions (p<0.0001; Fig. 1C, expanded scale in Supplemental Fig. S2B) relative to controls. Importantly, maximal taurine release in LRRC8A-null cells did not differ between the 30% and 50% osmotic reduction conditions (0.52% vs. 0.42%, respectively; p=0.512, Supplemental Fig. S2D). Because this residual release mechanism in bLRRC8A KO astrocytes is insensitive to the degree of cell swelling, our findings support activation of a VRAC-independent process and argue against incomplete Cre recombination or cell lysis.

### LRRC8A-containing VRACs dominate swelling-induced taurine release in the brain and regulate tissue taurine content

Having established a predominant role for LRRC8A-containing VRACs in hypoosmotic taurine release from cultured astrocytes, we next determined whether this pathway also mediates swelling-induced taurine release in the brain *in vivo*. To address this, we performed microdialysis experiments to measure extracellular taurine levels in the brain during perfusion with hypoosmotic aCSF (110 mosmol/kg H_2_O, ∼60% reduction). A larger reduction in medium osmolality was necessary to account for equilibration between the microdialysis probe and the surrounding extracellular fluid. Due to the relatively large size of the microdialysis probe (∼2-mm membrane length), we chose to restrict sampling to the striatum (∼20-37mm^3^) (57), which accommodated the full probe length. In the striatum, basal extracellular taurine levels were ∼6.2 µM in both control and bLRRC8A KO brains (p= 0.9863 vs. control, Fig. 1D and Supplemental Fig. S3A-B). In control brains, perfusion with hypoosmotic solution triggered a transient increase in extracellular taurine that reached a peak concentration of ∼57 µM within 10 minutes followed by partial recovery (Fig. 1D). In bLRRC8A KO brains, swelling-induced taurine release was nearly abolished with a ∼75% reduction in peak taurine levels (p=0.002, Fig. 1D) and a ∼92% decrease in integral taurine release under hypoosmotic conditions (p<0.0001, Fig. 1D). Together, these data show that swelling-induced taurine release in the brain is primarily mediated by VRACs, with minimal contribution from alternative pathways.

We previously reported (50) that bLRRC8A KO animals exhibit a 56% increase in brain tissue taurine levels as compared with littermate controls (p<0.0001, Fig. 1E). This finding is critical for interpreting our results and is therefore reproduced in Fig. 1E with permission. The increase in taurine content was restricted to the brain, as serum taurine levels measured from trunk blood were not significantly different between bLRRC8A KO animals and littermate controls (p=0.449, Fig. 1F). Together, these findings show that *Lrrc8a* deletion modifies brain taurine content without measurably altering systemic taurine levels.

### *Lrrc8d* is highly expressed in the mouse brain

While LRRC8A is obligatory for the formation of functional VRACs, the taurine permeability of these channels is influenced by their associated LRRC8 subunits. We therefore characterized the endogenous expression of all five *Lrrc8* transcripts in the mouse brain using publicly available RNA-seq datasets from the Human Protein Atlas (Fig. 2A). This analysis revealed substantial regional differences in the relative abundance of *Lrrc8a-d* transcripts. *Lrrc8e* expression was low to undetectable in all regions and was excluded from the presented heatmap. Across the mouse brain regions examined, *Lrrc8b* and *Lrrc8d* were the most abundant transcripts, although their levels varied considerably among regions (Fig. 2A). We next focused on three select regions—cerebral cortex, striatum, and hypothalamus—to explore *Lrrc8* transcript expression in a quantitative manner. In the cerebral cortex, the relative abundance of *Lrrc8* mRNA followed the pattern *Lrrc8b* > *Lrrc8d* ≈ *Lrrc8a* > *Lrrc8c* (Fig. 2C). The striatum, which was the site of microdialysis in our *in vivo* experiments, showed a similar expression profile (Fig. 2D). In contrast, the mouse hypothalamus exhibited a distinct shift in *Lrrc8* transcript distribution, with *Lrrc8d* emerging as the most abundant transcript, followed by *Lrrc8a* (Fig. 2E). Relative to the cortex, hypothalamic *Lrrc8b* expression was reduced by ∼62%, whereas *Lrrc8a* and *Lrrc8d* expression were increased by ∼2.2-fold and ∼1.6-fold, respectively (compare Fig. 2C and 2E).

**Figure 2:**
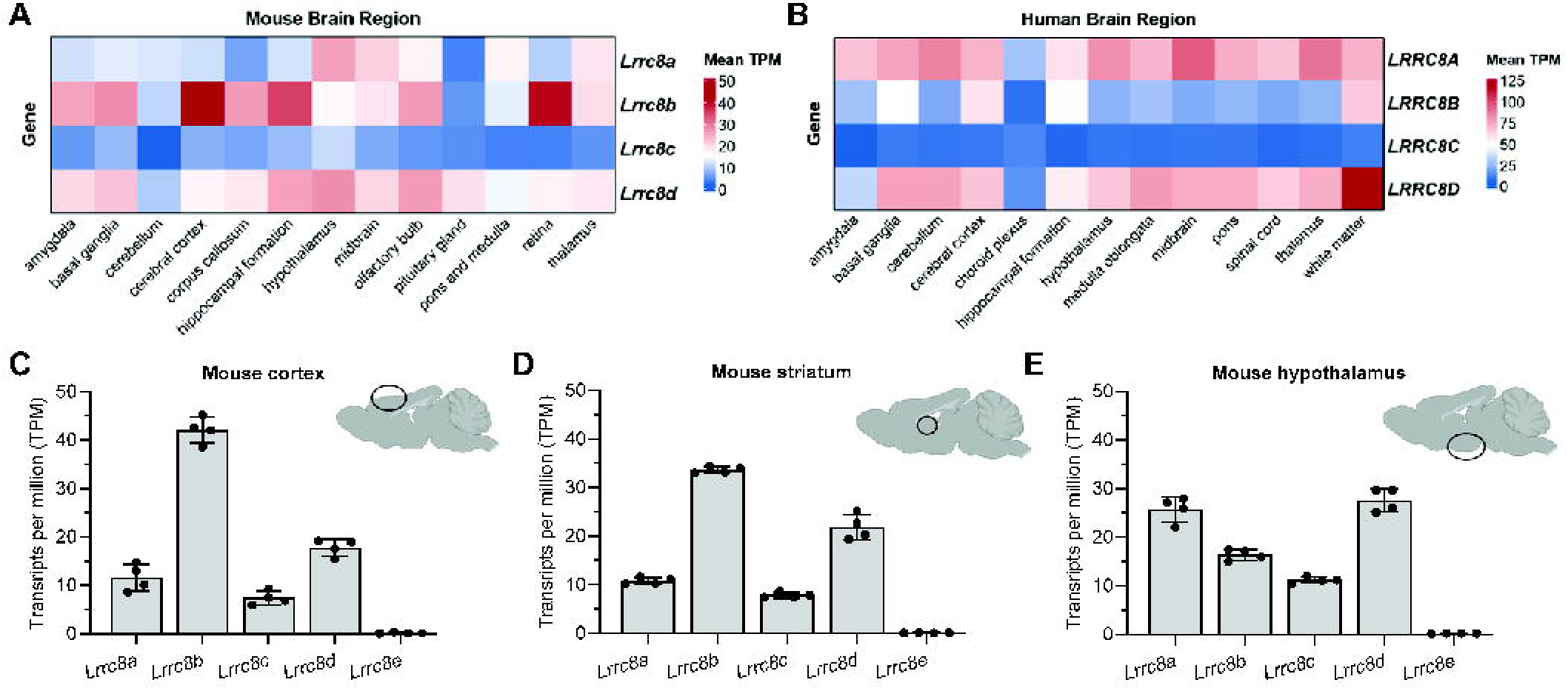
Region-specific analysis of VRAC subunit expression across the mouse and human brain. A, Heatmap depicting *Lrrc8a-d* transcript abundance in thirteen mouse brain regions. Values represent the mean transcripts per million (TPM) from four mice of both sexes. Data were downloaded from the Human Protein Atlas on April 29, 2026. **B**, Heatmap depicting *LRRC8A-D* transcript abundance in thirteen human brain regions. Values represent the mean TPM from samples obtained from the Human Protein Atlas on September 16, 2026. The dataset comprises 966 samples, which were grouped into thirteen major brain regions for this analysis. **C-E**, Quantification of *Lrrc8a-e* transcript abundance in the mouse cerebral cortex (**C**), striatum (**D**), and hypothalamus (**E**) using the samples analyzed in panel *A*.

To determine whether the expression pattern of VRAC subunits is conserved across species, we analyzed publicly available RNA-seq data in the human CNS using the Human Protein Atlas. In contrast to the mouse brain, *LRRC8A* was the most abundant transcript across the human brain regions examined, followed by *LRRC8D* (Fig. 2B). *LRRC8E* expression was again low to undetectable across all regions and was excluded from the presented heatmap. We performed a similar analysis using human RNA-seq datasets from the Genotype-Tissue Expression (GTEx) Project datasets and established a nearly identical relative abundance pattern (Supplemental Fig. S4). Together these findings show that *Lrrc8d* or *LRRC8D* was among the top two most abundant subunit transcripts in both species.

### Validation of siRNA-mediated knockdown of LRRC8C and LRRC8D

Previous studies have identified LRRC8D as an important determinant of taurine permeability; however, LRRC8C may also contribute (13, 29, 31, 33). Here, we explored the individual contributions of these two VRAC subunits to endogenous taurine-permeable VRACs in wild type mouse astrocytes using an RNAi approach and a radiotracer release assay.

The knockdown efficacy and target specificity of all siRNA constructs used was validated by qRT-PCR and western blotting in our recent work (32). To confirm that high siRNA efficacy is preserved in the present experiments, we performed additional western blot analysis in siRNA-treated astrocytes. Consistent with previous findings, the *Lrrc8c*-targeting siRNA (siC1) reduced LRRC8C immunoreactivity by ∼90% compared with cells treated with a negative-control siRNA (siNC; p=0.002; Fig. 3A-B; full Western blot dataset in Supplemental Fig. S5). Similarly, transfection with the *Lrrc8d*-targeting siRNA (siD4) reduced LRRC8D immunoreactivity by ∼92% compared with siNC-treated cells (p<0.0001; Fig. 3C-D; full Western blot dataset in Supplemental Fig. S6). These findings confirm robust protein-level depletion of the targeted LRRC8 subunits.

**Figure 3:**
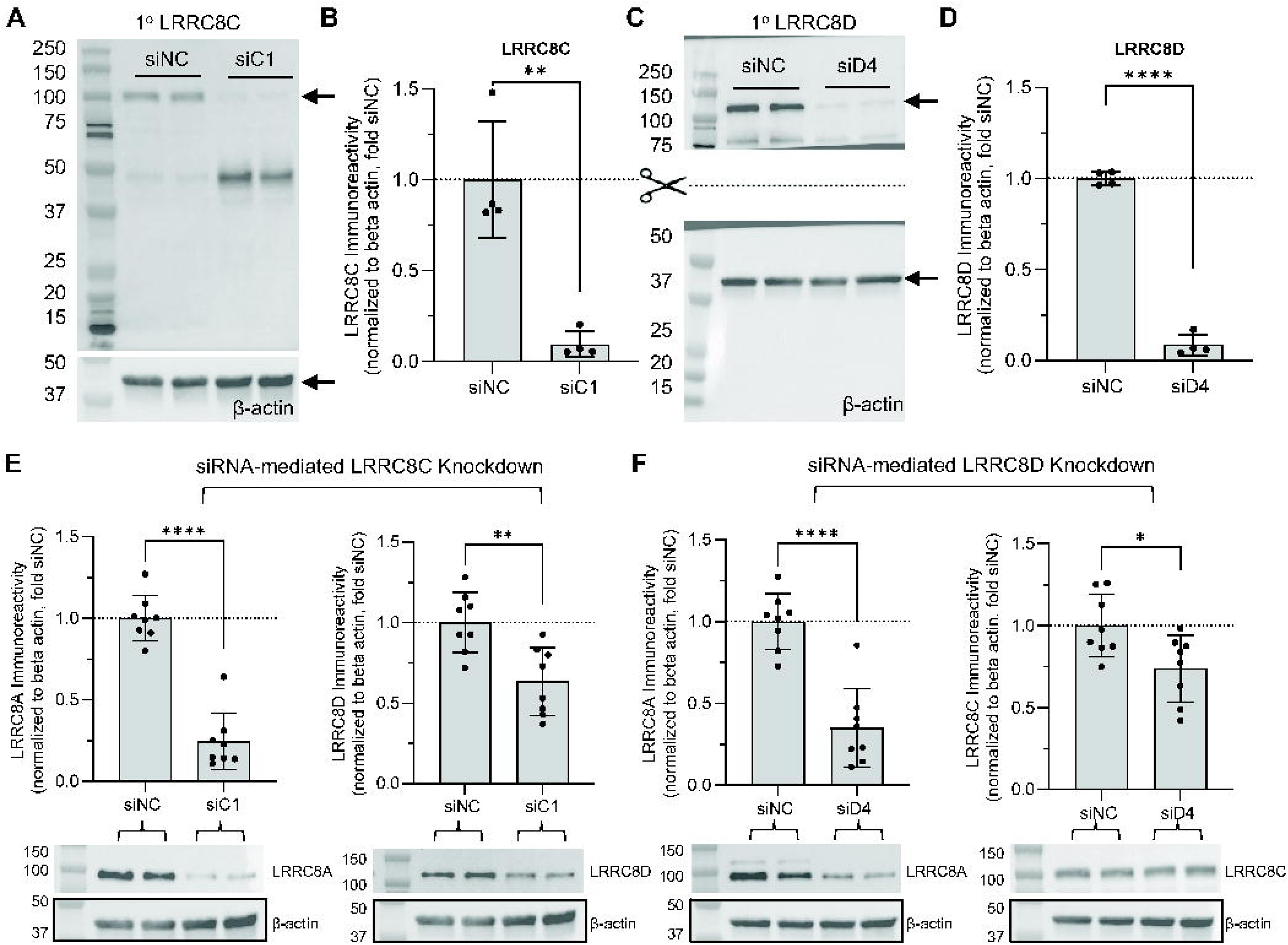
Subunit-specific LRRC8 knockdowns alter the protein levels of partner LRRC8 subunits. **A**, Representative western blot of LRRC8C immunoreactivity and corresponding β-actin signal from protein lysates prepared 96h post siRNA transfection with either negative-control siRNA siNC or the *Lrrc8c*-targeting siC1. **B**, Quantification of LRRC8C protein levels normalized to β-actin and presented relative to the average signal from siNC samples. Data are the mean values ± SD from four independent cultures per group. **p<0.01, unpaired t-test. **C,** Representative western blot of LRRC8D immunoreactivity and corresponding β-actin signal from protein lysates prepared 96h post siRNA transfection with either siNC or the *Lrrc8d*-targeting siD4. **D**, Quantification of LRRC8D protein levels normalized to β-actin and presented relative to the average signal from siNC samples. Data are the mean values ± SD from four independent cultures per group. ****p<0.0001, unpaired t-test. **E**, Top: Quantification of LRRC8A and LRRC8D protein levels in astrocytes treated with siNC or siC1, normalized to β-actin and expressed relative to siNC. Data are the mean values ± SD from eight independent cultures per group. ****p<0.0001, **p<0.01; unpaired t-test. Bottom: Representative western blot of LRRC8A or LRRC8D immunoreactivity and matching β-actin signal. LRRC8A and LRRC8D were probed on the same membrane, therefore, the beta-actin image is the same for both panels. **F**, Top: Quantification of LRRC8A and LRRC8C protein levels in astrocytes treated with siNC or siD4, normalized to β-actin and expressed relative to siNC. Data are the mean values ± SD from eight independent cultures per group. **** p<0.0001, *p<0.05; unpaired t-test. Bottom: Representative western blot of LRRC8A or LRRC8C immunoreactivity and matching β-actin signal.

### LRRC8 depletion differentially alters the protein abundance of associated subunits

Based on our recent report that LRRC8 subunits mutually regulate one another’s protein stability (32), we examined whether depletion of LRRC8C or LRRC8D alters the abundance of partner LRRC8 subunits in the current preparations. Immunoreactive bands for LRRC8A, LRRC8C, and LRRC8D differed in apparent molecular weight (see representative blots in Supplemental Figs. S5-S8). Together with the subunit-specific effects of siRNA knockdown, these distinct band patterns support the specificity of each antibody [see extended discussion in (32)].

The siRNA-mediated knockdown of LRRC8C or LRRC8D resulted in limited reciprocal regulation of protein levels of these two LRRC8 subunits. LRRC8C knockdown reduced LRRC8D immunoreactivity by 37% (p=0.003, Fig. 3E, full Western blot dataset in Supplemental Fig. S7), whereas LRRC8D knockdown reduced LRRC8C immunoreactivity by 26% (p=0.019, Fig. 3F, full Western blot dataset in Supplemental Fig. S8). In contrast, depletion of either subunit had a strong impact on LRRC8A protein levels. Specifically, LRRC8C knockdown decreased LRRC8A signal by 75% (p<0.0001, Fig. 3E, full Western blot dataset in Supplemental Fig. S7), and LRRC8D knockdown decreased it by 65% (p<0.0001, Fig. 3F, full Western blot dataset in Supplemental Fig. S8). Because each siRNA selectively reduced the mRNA expression of its target subunit without altering the transcript levels of the other *Lrrc8* subunits (32), the observed reductions in partner subunit protein levels suggest altered protein stability. Together, these findings support substantial contributions of both LRRC8C and LRRC8D to LRRC8A stability, with more modest effects on each other’s protein abundance.

### LRRC8D-containing VRACs dominate swelling-induced taurine release

Having validated robust siRNA-mediated depletion of LRRC8C and LRRC8D, we next assessed their subunit-specific contributions to swelling-activated taurine release using [^3^H]taurine as a radiotracer. Under isotonic conditions, baseline taurine release from siNC-treated control cells was low, averaging ∼0.14% per minute of the total [^3^H]taurine content (Fig. 4A). In control cells, a 30% reduction in extracellular osmolality (to 200 mosmol/kgH_2_O) triggered a rapid increase in taurine release reaching a peak rate of 3.1% within 2 minutes (Fig. 4A). When release was integrated over the entire response period, total taurine efflux reached ∼15% of the cellular [^3^H]taurine content (Fig. 4Aii).

**Figure 4:**
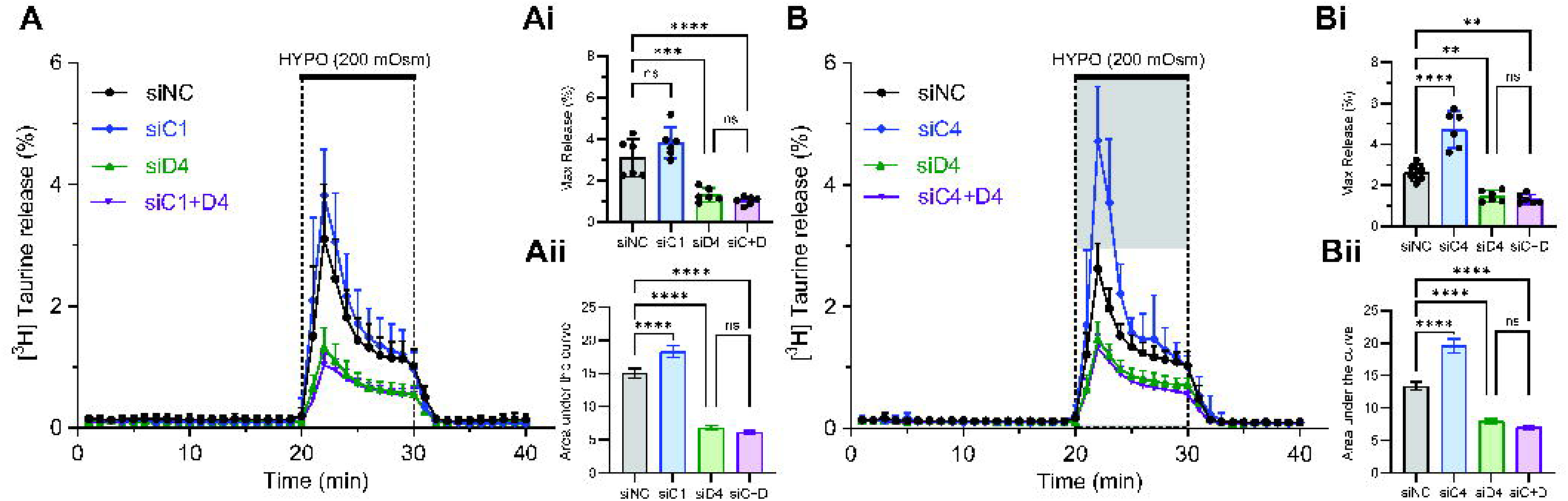
LRRC8D-containing VRACs dominate swelling-activated taurine release. **A**, Kinetics of VRAC-mediated [^3^H]taurine release from astrocyte cultures treated with siNC, siC1, siD4, or siC1+siD4 and exposed to a 30% reduction in medium osmolality. Data are the mean values ± SD in six independent transfections in two independently prepared cell cultures. *Insets*: Statistical comparison of the maximal (*Ai*) and integral (*Aii*) [^3^H]taurine release values in panel *A*. ****p<0.0001, ***p<0.001, one-way ANOVA with Tukey’s multiple comparisons test. **B**, Kinetics of VRAC-mediated [^3^H]taurine release from astrocyte cultures treated with siNC, siC4, siD4, or siC4+siD4 and exposed to a 30% reduction in medium osmolality. Data are the mean values ± SD in six independent transfections in two independently prepared cell cultures. *Insets*: Statistical comparison of the maximal (*Bi*) and integral (*Bii*) [^3^H]taurine release values in panel *B*. ****p<0.0001, **p<0.01, one-way ANOVA with Tukey’s multiple comparisons test.

Unexpectedly, knockdown of LRRC8C produced a paradoxical potentiation of [^3^H]taurine efflux, with a non-significant ∼22% increase in peak taurine release (p=0.211, Fig. 4Ai) and a significant ∼22% increase in integral taurine efflux under hypoosmotic conditions (p<0.0001, Fig. 4Aii). In contrast, LRRC8D knockdown strongly reduced swelling-induced taurine release, with a >58% reduction in maximum release (p=0.0004, Fig. 4Ai) and a ∼55% decrease in integral release under hypoosmotic conditions relative to controls (p<0.0001, Fig. 4Aii).

To determine whether LRRC8C and LRRC8D function redundantly within the same VRAC complexes, we performed additivity experiments by simultaneously knocking down both subunits. We previously confirmed that co-transfection with siC1 and siD4 reduced target gene expression by ≥80%, both individually and in combination (32). Combined knockdown of LRRC8C and LRRC8D did not further reduce swelling-activated taurine release beyond that observed with LRRC8D knockdown alone (p= 0.859 siC1+siD4 vs. siD4, Fig. 4A), indicating no additive or synergistic effect.

To further investigate the unexpected increase in [^3^H]taurine efflux following LRRC8C knockdown and rule out off-target siRNA effects, we repeated the experiments using a different *Lrrc8c*-targeting siRNA construct (siC4) (Fig. 4B). Strikingly, knockdown of LRRC8C using siC4 resulted in an ≥80% increase in maximal taurine release (p<0.0001, Fig. 4Bi) and ∼46% increase in integral taurine release under hypoosmotic conditions (p<0.0001, Fig. 4Bii). In this set of experiments, LRRC8D knockdown again produced a robust >46% reduction in peak swelling-induced taurine release (p=0.006, Fig. 4Bi) and a ∼40% decrease in integral radiotracer release compared with control cells (p<0.0001, Fig. 4Bii). These results show that the enhancement of VRAC-mediated taurine release following LRRC8C depletion is reproducible and unlikely to be due to an off-target effect of the siRNA.

### Gene-trap mutagenesis of LRRC8D results in incomplete loss of LRRC8D protein

Given the prominent role of the LRRC8D subunit in swelling-activated taurine release, we next utilized a commercially available “gene-trapped” *Lrrc8d* knockout mouse strain to examine the effects of *Lrrc8d* deletion *in vivo*. Crossing heterozygous animals yielded wild-type (WT), heterozygous (HET), and knockout (KO) offspring at frequencies close to the expected Mendelian ratio (1:2:1, Fig. 5A), indicating normal viability of *Lrrc8d*-deficient mice. Analysis of body weight at 8-10 weeks of age revealed a main effect of sex (p<0.0001, Fig. 5B), but no significant effect of genotype (p=0.302, Fig. 5B).

**Figure 5:**
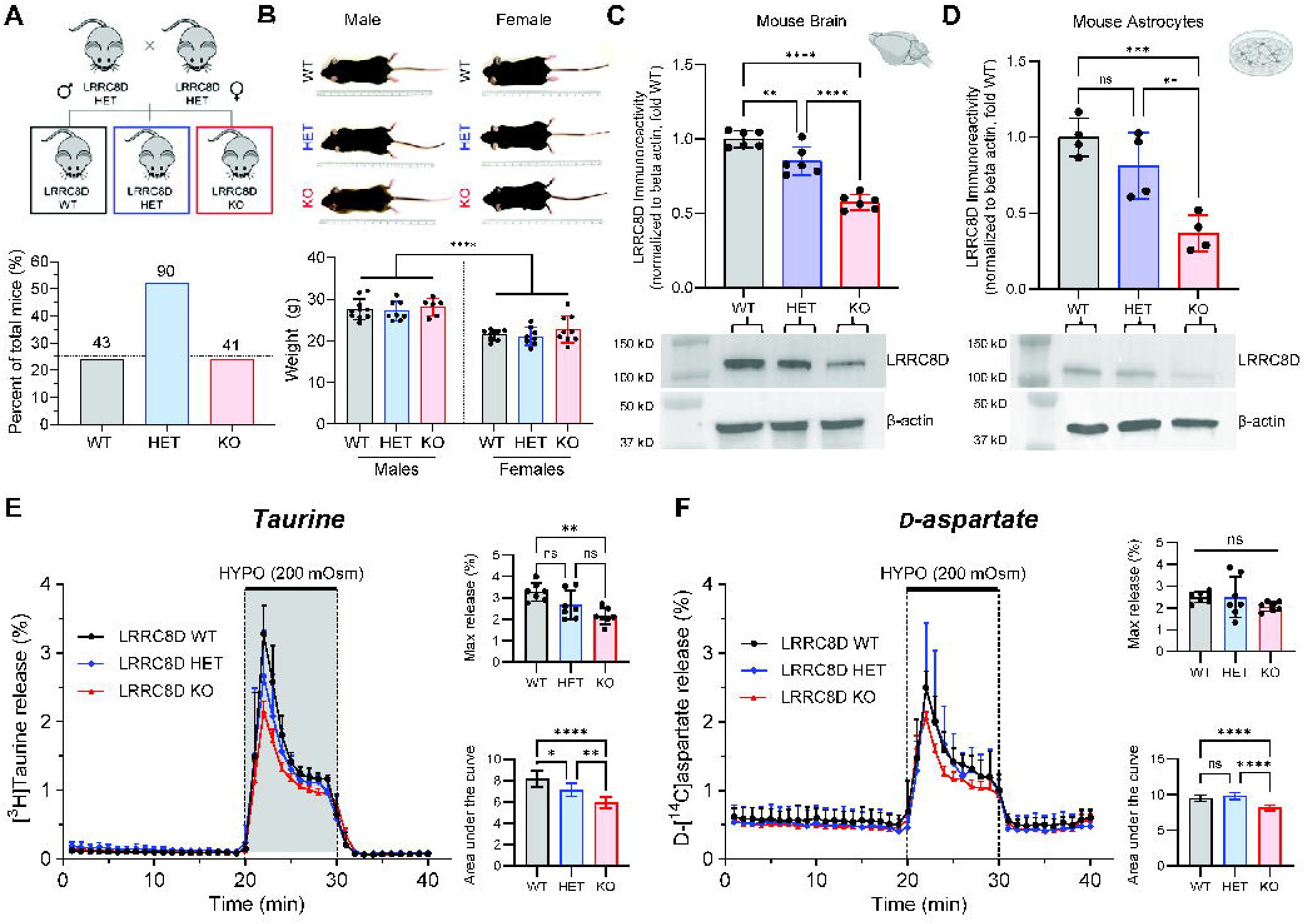
Gene-trap mutagenesis of LRRC8D results in a partial reduction in LRRC8D expression and mild decrease in VRAC-mediated taurine release. **A**, Breeding strategy for genetic deletion of LRRC8D using gene-trap mutagenesis. The three resulting genotypes are: *Lrrc8d* (WT), *Lrrc8d* (HET), and *Lrrc8d* (KO) with the expected Mendelian ratio of 1:2:1. Numbers above bars indicate the total number of live-born animals from each genotype. **B**, Representative images and body weight quantification for males and females from each genotype. Body weights of animals of both sexes measured at 8-10 weeks. Data are the mean values ± SD in 6-10 animals per group. ****p<0.0001 (effect of sex), two-way ANOVA with Šídák’s multiple comparisons test. **C**, Western blot analysis of LRRC8D protein levels in forebrain lysates, normalized to β-actin of each sample and additionally to WT controls in each experiment. Data are the mean values ± SD from six different animals per genotype. ****p<0.0001, **p<0.01, one-way ANOVA with Tukey’s multiple comparisons test. **D**, Western blot analysis of LRRC8D protein levels in primary astrocyte cultures, normalized to β-actin of each sample and additionally to WT controls in each set. Data are the mean values ± SD from four different cultures per genotype. ***p<0.001, **p<0.01, one-way ANOVA with Tukey’s multiple comparisons test. **E**, Kinetics of VRAC-mediated [^3^H]taurine release from astrocyte cultures from control, *Lrrc8d* HET, and *Lrrc8d* KO genotypes exposed to a 30% reduction in medium osmolality. Data are the mean values ± SD in seven independent experiments per genotype in three different astrocyte cultures. *Insets:* Statistical comparison of the maximal (top) and integral (bottom) [^3^H]taurine release values in panel *E*. ****p<0.0001, **p<0.01, *p<0.05; one-way ANOVA with Tukey’s multiple comparisons test. **F**, Kinetics of VRAC-mediated D-[^14^C]aspartate release from astrocyte cultures from control, *Lrrc8d* HET, and *Lrrc8d* KO genotypes exposed to a 30% reduction in medium osmolality. Data are the mean values ± SD in seven independent experiments per genotype in three different astrocyte cultures. *Insets*: Statistical comparison of the maximal (top) and integral (bottom) D-[^14^C]aspartate release values in panel *F*. ****p<0.0001, one-way ANOVA with Tukey’s multiple comparisons test.

To confirm effective depletion of LRRC8D protein, we performed western blot analysis on forebrain protein lysates. Relative to WT brains, LRRC8D immunoreactivity was reduced by ∼15% in HET mice (p=0.006, Fig. 5C) and by ∼42% in KO mice (p<0.0001, Fig. 5C, full Western blot dataset in Supplemental Fig. S9), indicating that this gene-trapped strategy produces only partial loss of LRRC8D protein. Consistent with these findings, primary astrocyte cultures derived from these animals showed a non-significant ∼19% reduction of LRRC8D protein in HET cells (p=0.277, Fig. 5D) and a partial ∼63% reduction in KO astrocytes (p=0.001, Fig. 5D, full Western blot dataset in Supplemental Fig. S10). While this “gene trapped” model was supposed to result in a constitutive knockout, we found only partial depletion of the LRRC8D protein.

We next determined if this partial LRRC8D depletion alters swelling-activated taurine release. To further establish whether any observed effects were specific for taurine, primary cultured astrocytes from WT, HET, and LRRC8D-deficient mice were simultaneously loaded with [^3^H]taurine and D-[^14^C]aspartate, a non-metabolizable analogue of glutamate, allowing simultaneous measurement of swelling-induced efflux of both amino acids in the same astrocyte cultures.

Under isotonic conditions, basal [^3^H]taurine release was low and did not differ between genotypes, averaging ∼0.1% per minute of total tracer content (Fig. 5E). Exposure to hypoosmotic medium induced a rapid increase in taurine efflux in WT astrocytes, reaching a peak release rate of 3.3% within 2 min (Fig. 5E). Heterozygous deletion of LRRC8D produced a mild phenotype, with a non-significant ∼18% reduction in peak swelling-induced taurine release (p=0.084, Fig. 5E) and a small but significant ∼12% decrease in total integrated taurine efflux under hypoosmotic conditions relative to WT controls (p=0.023, Fig. 5E). In comparison, LRRC8D KO cells exhibited a more pronounced impairment, with a ∼34% reduction in peak taurine release (p=0.002, Fig. 5E) and an ∼27% decrease in total hypoosmotic taurine efflux compared with WT cells (p<0.0001, Fig. 5E). These findings demonstrate that even partial depletion of LRRC8D (∼63% reduction in protein expression) attenuates swelling-evoked taurine efflux, supporting an important role for this VRAC subunit in conferring taurine permeability to astrocytic VRACs.

In the same cells, basal D-[^14^C]aspartate release under isotonic conditions averaged ∼0.5% per minute of total tracer content in all three genotypes, 5-fold higher than the observed basal [^3^H]taurine release (compare Fig. 5F and Fig. 5E). Hypoosmotic stimulation triggered robust D-[^14^C]aspartate efflux in WT astrocytes, with release reaching a peak rate of ∼2.5% within 2 min (Fig. 5F). In contrast to taurine, LRRC8D depletion had only a minor impact on glutamate release. One-way ANOVA revealed no significant effect of genotype on maximal D-[^14^C]aspartate efflux (p=0.292, Fig. 5F). LRRC8D HET astrocytes showed no detectable changes in either peak or integral release values (Fig. 5F). LRRC8D KO astrocytes exhibited a small but statistically significant 14% decrease in total integrated efflux (p<0.0001, Fig. 5F). Overall, these findings show that LRRC8D depletion has a greater impact on taurine release than glutamate release.

### LRRC8D depletion does not impair systemic osmoregulation following acute hyper-osmotic challenge

Given the proposed role of astrocyte-derived taurine in regulating hypothalamic osmoregulatory circuits, we next investigated whether impaired LRRC8D-dependent taurine release affects systemic fluid and electrolyte homeostasis. First, we tested the ability WT, LRRC8D HET, and LRRC8D-deficient mice to regulate their osmolality following a hyperosmotic challenge. Acute elevation of plasma osmolality is expected to activate magnocellular neurosecretory neurons in the SON and PVN and promote AVP release (Fig. 6A). Mice received an i.p. injection of 2M NaCl and we measured serum osmolality over time.

**Figure 6:**
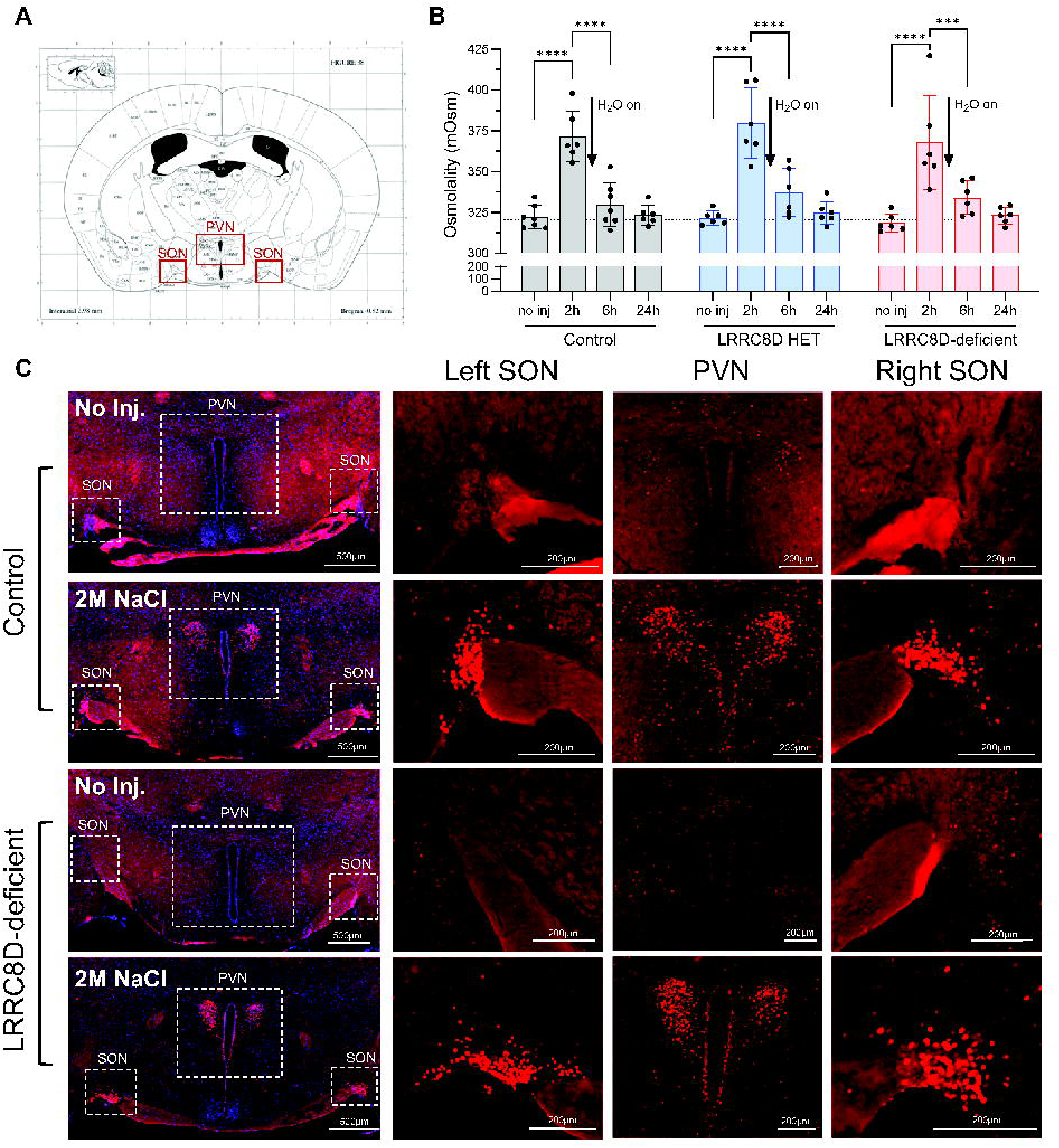
Depletion of LRRC8D does not impair systemic osmoregulation in response to acute hyperosmotic challenge. **A**, Schematic representation of the location of the SON and PVN based on the Paxinos and Franklin mouse atlas coordinates. **B**, Effect of acute hyperosmotic challenge on serum osmolality in mice with gene-trap deletion of *Lrrc8d*. Anesthetized animals were injected i.p. with 2 M NaCl at time 0 and their trunk blood was collected at specified times. Data are the mean values ± SD in 6-7 animals per group per time. ****p<0.0001, ***p<0.001; two-way ANOVA with Tukey’s multiple comparisons test. **C**, Representative images of cFos immunoreactivity (red) and DAPI nuclear staining (blue) in the SON and PVN of WT and LRRC8D-deficient animals under baseline conditions (no inj.) and 2 h after i.p. injection of 2 M NaCl. *Left*, low-magnification images of the SON and PVN. Scale bar = 500 µm. *Right*, higher-magnification images of the selected regions of interest. Scale bar = 200 µm.

Baseline serum osmolality was similar among WT, HET, and LRRC8D-deficient mice, averaging 322, 321, and 318 mosmol/kg H_2_O, respectively (Fig. 6B). I.p. injection of 2 M NaCl produced a rapid increase in serum osmolality that peaked at ∼373 mosmol/kg H_2_O during 2 h water restriction, corresponding to an approximately 17% increase from baseline. Following restoration of water access, serum osmolality progressively declined and returned to baseline within 24 h in all three genotypes (Fig. 6B). Two-way ANOVA revealed a significant main effect of time (p<0.0001, Fig. 6A) but no effect of genotype (p=0.406, Fig. 6B). These findings indicate that partial LRRC8D depletion does not impair recovery from an acute hyperosmotic challenge.

In parallel experiments, we confirmed activation of magnocellular neurons in the SON and PVN following the hyperosmotic challenge by labeling cFos, a widely used marker of neuronal activity (58). At 2 h after NaCl administration (maximal increase in serum osmolality), we found a robust increase in cFos expression in both the PVN and SON (Fig. 6C). Qualitative examination of the representative sections revealed no apparent differences in cFos labeling between WT and LRRC8D-deficient mice (Fig. 6C). Because systemic osmoregulatory responses did not differ between genotypes, we did not pursue quantitative analysis of cFos labeling in the SON or PVN.

### LRRC8D depletion or LRRC8A KO do not impair systemic osmoregulation following acute water intoxication

We next examined whether LRRC8D depletion impairs recovery from acute water intoxication. Hypoosmotic conditions induce cellular swelling and activate VRACs, allowing us to assess the physiological consequences of impaired VRAC-mediated taurine release in genetically modified animals. Hypoosmolality rapidly inhibits magnocellular neurons in the SON and PVN, thereby reducing pituitary AVP release, a response thought to be mediated in part by taurine signaling (45, 46). In rodents, circulating AVP has a half-life of 1-8 min, allowing animals to rapidly transition to regulatory diuresis (59).

During hypoosmolality, taurine release may help inhibit magnocellular neurons in the SON and PVN, thereby reducing pituitary AVP secretion.

Mice received an i.p. injection of sterile water equivalent to 7.5% of body weight. Serum osmolality decreased by approximately 12%, reaching on average 282 mosmol/kg H_2_O within 30 min, and then recovered toward baseline over approximately 2 h in all three genotypes (Fig. 7A). The same baseline serum osmolality values from non-injected animals were used for both the hyperosmotic and hypoosmotic challenges. Two-way ANOVA revealed a significant effect of time (p<0.0001, Fig. 7A) but no significant effect of genotype (p=0.787, Fig. 7A). Together with the hyperosmotic challenge results, these findings indicate that LRRC8D deficiency does not measurably impair systemic osmotic recovery following either acute challenge.

**Figure 7:**
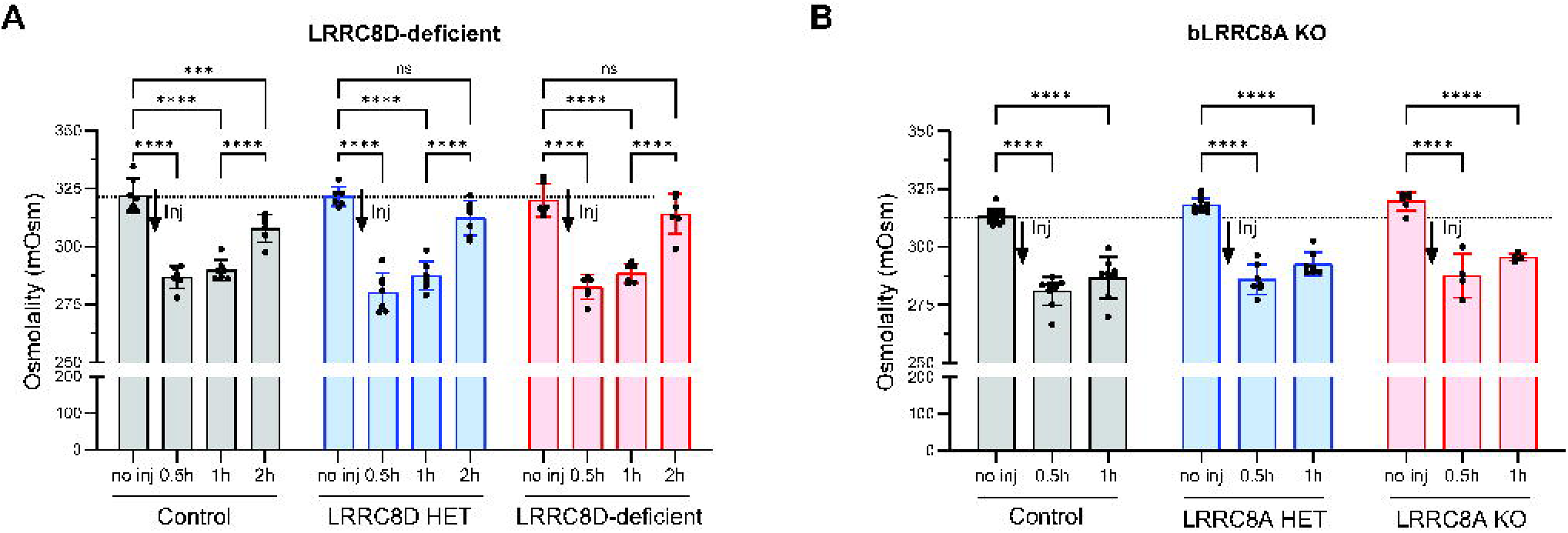
Loss of LRRC8A or LRRC8D does not impair recovery from acute water intoxication. **A**, Effect of acute systemic hypoosmotic challenge on serum osmolality in gene-trapped *Lrrc8d* mice. Anesthetized animals were injected i.p. with sterile water (7.5% body weight) at time 0 and their trunk blood was collected at specified times. Data are the mean values ± SD in 6-7 animals per group per time. ****p<0.0001, ***p<0.001; two-way ANOVA with Tukey’s multiple comparisons test. **B**, Effect of acute systemic hypoosmotic challenge on serum osmolality in mice with brain-wide bLRRC8A KO. Control (fl/+ and fl/fl), LRRC8A HET, and bLRRC8A KO mice were injected i.p. with sterile water at time 0 and their trunk blood was collected at specified times. Data are the mean values ± SD in 4-12 animals per group per time. ****p<0.0001; two-way ANOVA with Tukey’s multiple comparisons test.

Because the LRRC8D gene-trap model retains partial LRRC8D protein expression, residual protein may be sufficient to preserve VRAC function *in vivo*. We therefore repeated the acute hypoosmotic challenge in bLRRC8A KO mice, which lack the obligatory LRRC8A subunit in the brain and consequently lack functional VRACs in neurons, astrocytes, and oligodendrocytes. *Lrrc8a*^fl/+^ and *Lrrc8a*^fl/fl^ animals were combined as controls. Given the rapid recovery observed in the LRRC8D-deficient cohort, we shortened the observation period from 2 h to 1 h.

Baseline serum osmolality was similar among control, LRRC8A HET, and bLRRC8A KO mice, averaging 313, 318, and 320 mosmol/kg H_2_O, respectively (Fig. 7B). Thirty minutes after i.p. injection of sterile water, serum osmolality fell to ∼280-285 mosmol/kg H_2_O in all three genotypes, representing an approximately 10% reduction from baseline (Fig. 7B). Two-way ANOVA revealed significant main effects of time (p<0.0001, Fig. 7B) and genotype (p=0.0002, Fig. 7B), but no significant time × genotype interaction (p=0.9867, Fig. 7B). Thus, although serum osmolality differed overall among genotypes, we detected no genotype-dependent differences in the time course of the response to the hypoosmotic challenge.

## DISCUSSION

In this study, we determined the LRRC8 subunit requirements for taurine efflux through endogenous brain VRACs and examined whether VRAC-mediated taurine release contributes to systemic osmoregulation. Our findings support three main conclusions. First, LRRC8A-containing VRACs provide the primary pathway for swelling-induced taurine release in the mouse brain. Second, LRRC8D is critical for conferring taurine permeability of VRACs. Third, despite LRRC8-containing VRACs central role in swelling-induced taurine release, they appear to be dispensable for systemic osmoregulation in response to acute osmotic challenges.

A major finding of this study is that LRRC8A-containing VRACs constitute the primary endogenous pathway for swelling-activated taurine release in the brain. Hypoosmotic stimulation has long been known to increase extracellular levels of taurine and other amino acids in the brain (60–64). Although pharmacological studies implicated VRACs in this process (48, 51, 64, 65), their unknown molecular identity precluded direct genetic validation. The identification of LRRC8 proteins as VRAC components enabled genetic studies confirming their contribution to swelling-activated taurine efflux in model cell lines and primary astrocyte cultures (12, 13, 20, 21, 31). Extending these findings *in vivo*, we found that LRRC8A deletion largely abolished swelling-induced taurine release in brain microdialysis experiments. Interestingly, a small residual release of taurine was observed under hypoosmotic conditions both *in vivo* and in cultured astrocytes. This residual release is unlikely to reflect cell lysis or incomplete *Lrrc8a* deletion, as it was not further increased by a greater reduction in extracellular osmolarity (i.e., it was insensitive to the degree of cell swelling). Instead, the residual response likely reflects a minor contribution from an alternative swelling-stimulated release pathway. Several mechanisms have been proposed to mediate volume-sensitive organic osmolyte release, including Maxi-Cl⁻ channels (66, 67), Tweety homolog proteins TTYH1-3 (68), Ca^2+^-activated BEST1 channels (69, 70), and possibly connexin 43 hemichannels (71). Although the molecular identity of the residual release pathway remains unresolved, its quantitative contribution is minor.

Beyond VRACs role in swelling-activated taurine release, these channels appear to contribute to basal taurine efflux and brain taurine homeostasis. We previously reported that total brain taurine content was 56% higher in bLRRC8A KO mice than in littermate controls (50). Because *Nestin*^Cre^-mediated deletion is restricted to a subset of CNS cells, the increase in intracellular taurine within targeted cells may be greater than the increase measured in whole-brain tissue. We speculate that, after being taken inside cells by the Na^+^-dependent TauT, taurine becomes trapped following the loss of the VRAC-mediated efflux pathway. Supporting a role for VRACs in basal taurine efflux, we found that LRRC8A deletion reduced unstimulated taurine release from cultured mouse astrocytes, and similar findings were reported in rat astroglial cells (12, 31). In agreement with the idea that VRAC activity influences basal levels of extracellular neurotransmitters, Yang and colleagues found that astrocytic LRRC8A-containing VRACs mediate tonic release of GABA which contributes to cocaine-related behavioral responses in mice (72). Yet, despite reduced basal taurine release in cultured astrocytes, our microdialysis experiments detected no clear differences in baseline extracellular taurine levels between bLRRC8A KO mice and littermate controls. Baseline extracellular GABA was below the HPLC detection limit (data not shown). Preserved extracellular taurine levels could reflect compensatory release through alternative pathways, such as reverse operation of TauT facilitated by elevated intracellular taurine content. Overall, our findings indicate that LRRC8A deletion significantly alters taurine homeostasis in the brain.

Consistent with previous work in model cell lines (13, 33), our findings identify LRRC8D as a critical determinant of taurine conductance through endogenous VRACs in mouse astrocytes. The preferential permeability to taurine is likely based on its charge, as other net-neutral osmolytes, such as myo-inositol and GABA are released via the same pathway (13, 31). LRRC8A/LRRC8D-containing VRACs also provide a major uptake pathway for the chemotherapeutic agent cisplatin and the antibiotic blasticidin S (33, 73). This unique substrate specificity is likely determined by LRRC8D’s contribution to channel structure. Cryo-electron microscopy studies show that incorporation of LRRC8D widens the VRAC pore and increases the hydrophobicity of its selectivity filter (27, 74, 75). The physiological relevance of LRRC8D-containing VRACs in the brain is supported by the high abundance of *Lrrc8d* transcripts across brain regions, with some regional variation. In comparison, *Lrrc8c* expression is on average three times lower than *Lrrc8d*. This is potentially significant because LRRC8C-containing VRACs mediate glutamate release (13, 31, 32). *Lrrc8b* is also highly expressed in the mouse brain, although its functional contributions remain unclear. These expression patterns are consistent with our recently published qRT-PCR data from the mouse forebrain (32). Together with our functional findings, these observations suggest that taurine-permeable LRRC8A/LRRC8D-containing VRACs may play a dominant role in brain physiology.

In contrast to LRRC8D, LRRC8C was dispensable for swelling-activated taurine release. Unexpectedly, *Lrrc8c* knockdown enhanced VRAC-mediated taurine release, an effect reproduced with two independent *Lrrc8c*-targeting siRNA constructs. It is important to note that this potentiation occurred despite a nearly four-fold reduction in LRRC8A protein levels. There are two potential explanations of how LRRC8C can limit VRAC-mediated taurine release. First, LRRC8C and LRRC8D may compete for the obligatory LRRC8A subunit to form distinct channel populations with different substrate selectivities. In this model, LRRC8C depletion would favor the assembly of taurine-permeable LRRC8A/LRRC8D channels. Alternatively, LRRC8C may assemble with LRRC8A and LRRC8D to form channels with lower taurine permeability than LRRC8A/LRRC8D heteromers. Lutter et al. showed that co-expression of LRRC8C with LRRC8A and LRRC8D reduced the relative permeability to taurine compared with co-expression of LRRC8A and LRRC8D alone (13). Our recent work on glutamate-permeable VRACs (32) and earlier RNAi-based analysis of swelling-activated glutamate and taurine release in rat astrocytes (31) support the existence of distinct VRAC populations (LRRC8A/LRRC8C and LRRC8A/ LRRC8D) with different substrate selectivities.

In further support of the two-population model, reciprocal changes in LRRC8 subunit protein levels observed here and in our previous work (32) suggest that subunit stability depends on its assembly partners. Notably, changes in LRRC8C and LRRC8D have greater effects on LRRC8A protein levels than on each other. Because these changes occurred without corresponding changes in mRNA levels, they likely reflect altered protein stability (32). Consistent with these findings, Yu et al. recently reported that endothelial-specific deletion of *Lrrc8a* reduced LRRC8C protein abundance by >80%, while global *Lrrc8c* deletion reduced LRRC8A protein levels approximately four-fold (76). Although the two-population model requires further scrutiny, our findings show that LRRC8C is not required for taurine release and may limit the taurine permeability of endogenous astrocytic VRACs.

To assess the physiological relevance of LRRC8D-containing VRACs, we focused on the hypothalamus. Prior studies accumulated strong *ex vivo* evidence indicating that astrocytic taurine release modulates hypothalamic osmoregulatory circuits, with VRACs implicated as the most likely release pathway (45, 48, 49, 65). Nevertheless, VRACs’ contribution to hypothalamic function has not been definitively tested using tools of molecular genetics. Our finding of elevated hypothalamic expression of both *Lrrc8a* and *Lrrc8d* provided further rationale for examining the role of VRACs in osmoregulation. We initially tested commercially available mice with gene-trap-mediated disruption of *Lrrc8d* but detected no changes in systemic osmoregulation following either hyperosmotic or hypoosmotic challenges. Unfortunately, these results were inconclusive because we found only partial reductions in LRRC8D expression and VRAC-mediated taurine release in the gene-trap model. We therefore repeated the hypoosmotic challenge experiments in mice with brain-wide bLRRC8A KO, which abolishes brain LRRC8A expression and VRAC function ((50) and present work). Our current microdialysis experiments confirmed dramatic reductions in swelling-activated taurine release in bLRRC8A KO brains. Although we detected a main effect of genotype on serum osmolarity, the osmolarity response to water intoxication did not differ between control and LRRC8A-null mice. Thus, even under conditions expected to strongly activate brain VRACs, their deletion did not alter the systemic response to an acute hypoosmotic challenge.

The unexpected lack of an osmoregulation phenotype in the bLRRC8A KO has an important literature precedent. A recent study from the Jentsch laboratory examined the contributions of LRRC8A and LRRC8D to kidney function (77). Because global LRRC8A deletion is lethal, the authors used *Apoe*-Cre to target kidney cells, a strategy that also impacts brain glia (78). Conditional LRRC8A deletion caused renal injury, fibrosis, increased diuresis, and impaired water retention during water deprivation (77). Nevertheless, the circulating levels of arginine vasopressin were unchanged, suggesting unimpaired activity of the hypothalamic-pituitary axis (77). Global *Lrrc8d* deletion produced qualitatively similar renal injury and functional impairments (77). Our findings complement this work by directly measuring serum osmolality during water intoxication, a condition in which swelling-induced VRAC activation would be expected to contribute prominently. Several explanations, such as developmental compensation and/or substantial redundancy of the intrinsic osmosensory mechanisms (79) may account for these negative findings. Still, the combined observations in this study and that by López-Cayuqueo and colleagues (77) suggest that brain taurine-permeable VRACs are dispensable for hypothalamic osmoregulatory signaling.

In summary, our findings establish LRRC8A-containing VRACs as the primary pathway for swelling-activated taurine release in the mouse brain and identify LRRC8D as the main determinant of taurine permeability in mouse astrocytic VRACs. Notably, LRRC8A deletion increases total brain taurine content, supporting an important role for VRACs in taurine homeostasis. Together with the widespread expression of *Lrrc8d* transcripts in the brain, these findings suggest that LRRC8A/LRRC8D-containing VRACs may contribute to inhibitory taurine signaling in the CNS. Despite their central role in swelling-activated taurine release, these channels appear dispensable for hypothalamic control of systemic fluid and electrolyte homeostasis.

## DATA AVALIBILITY

All primary data included in this manuscript have been deposited in the Open Science Framework depository and can be accessed at https://doi.org/10.17605/OSF.IO/TSZNX. All original western blot images are available in figshare at https://doi.org/10.6084/m9.figshare.34625250.

## SUPPLEMENTAL MATERIAL

Supplemental Figs. S1-S10 can be found at https://doi.org/10.6084/m9.figshare.34625250.

## ACKNOWLEDGMENTS

The authors thank Dr. Alena Rudkouskya for help with LRRC8A western blot experiments and Dr. Jiayue Hong for contribution to pilot experiments exploring taurine release in LRRC8D KO astrocytes. Illustrations in Figs. 1, 2, 5, and 8 were created with BioRender.

## AUTHOR CONTRIBUTIONS

Conceived and designed research: M.L.C., J.W.N, P.D., and A.A.M. Performed experiments: M.L.C., J.W.N., P.D., and A.D.S. Analyzed data: M.L.C., J.W.N., P.D., A.D.S., A.L.S., and A.A.M. Interpreted results: M.L.C., J.W.N., P.D., and A.A.M. Prepared figures: M.L.C., J.W.N., and A.A.M. Drafted manuscript: M.L.C., J.W.N., and A.A.M. Edited and revised manuscript, M.L.C., J.W.N, P.D., A.D.S., A.L.S., and A.A.M. Approved final version of manuscript: M.L.C., J.W.N., P.D., A.D.S., A.L.S., and A.A.M.

