## Supplemental Figures S1-S10 for "LRRC8A/D-containing anion channels mediate swelling-activated taurine release in the mouse brain"

**Supplemental Figure S1**. The complete set of western blot images showing LRRC8A and β-actin immunoreactivity in four sets of protein lysates derived from LRRC8A fl/+, fl/fl, HET, and KO forebrains

**Supplemental Figure S2**. An extended analysis of basal and swelling-activated [$^3$H]taurine release in control and LRRC8A KO astrocytes from the datasets presented in main Figure 1B and 1C

**Supplemental Figure S3**. An extended analysis of taurine levels in microdialysate samples collected in control and bLRRC8A KO brains from the datasets presented in main Figure 1D

**Supplemental Figure S4.** Analysis of *LRRC8A-D* expression across 12 human brain regions

**Supplemental Figure S5**. The complete set of western blot images showing LRRC8C and β-actin immunoreactivity in four sets of protein lysates that were collected 96 hours post treatment with either negative control siRNA (siNC) or *Lrrc8c*-targeting siRNA (siC1)

**Supplemental Figure S6:** The complete set of western blot images showing LRRC8D and β-actin immunoreactivity in four sets of protein lysates that were collected 96 hours post treatment with either negative control siRNA (siNC) or *Lrrc8d*-targeting siRNA (siD4)

**Supplemental Figure S7**. The complete set of western blot images showing LRRC8A or LRRC8D expression in eight sets of protein lysates that were collected 96 hours post treatment with negative control siRNA (siNC) or *Lrrc8c*-targeting siRNA (siC1)

**Supplemental Figure S8**. The complete set of western blot images showing LRRC8A and LRRC8C immunoreactivity in eight sets of protein lysates that were collected 96 hours post treatment with negative control siRNA (siNC) or *Lrrc8d*-targeting siRNA (siD4)

**Supplemental Figure S9.** The complete set of western blot images showing LRRC8D and β-actin immunoreactivity in six sets of protein lysates derived from LRRC8D WT, HET, and KO forebrains

**Supplemental Figure S10.** The complete set of western blot images showing LRRC8D and β-actin immunoreactivity in four sets of protein lysates prepared from astrocyte cultures derived from LRRC8D WT, HET, and KO neonates

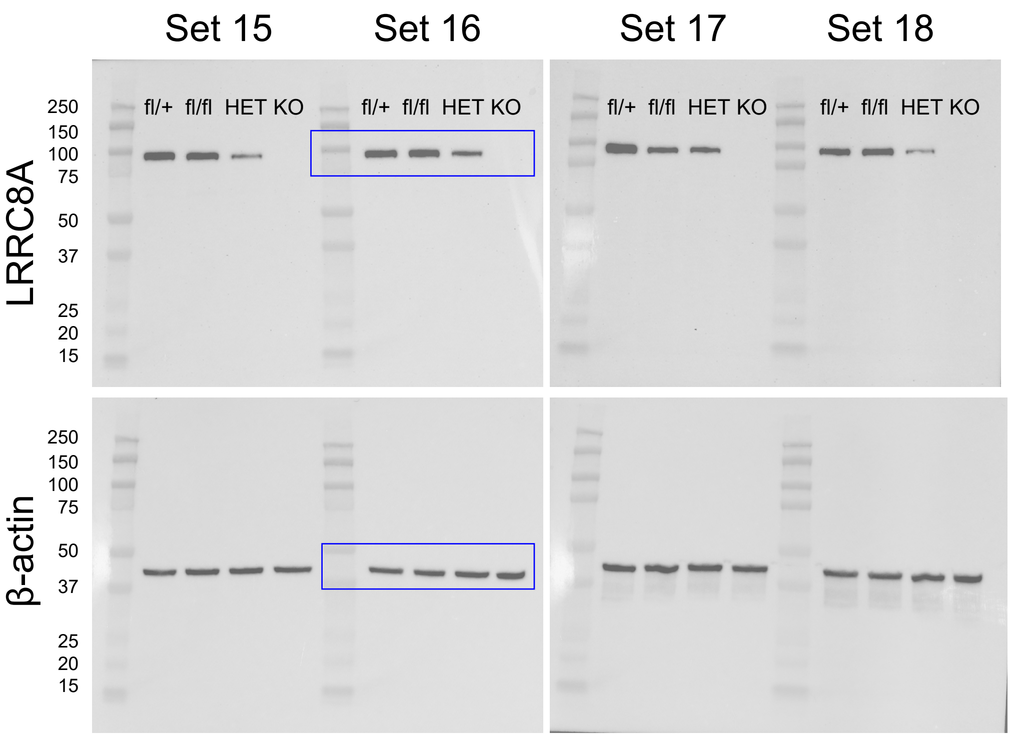


**The complete set of western blot images showing LRRC8A and β-actin immunoreactivity in four sets of protein lysates derived from LRRC8A fl/+, fl/fl, HET, and KO forebrains**

This is the full dataset of western blot images represented and quantified in the main Figure 1A. *Blue boxes* show the representative cropped images shown in main Figure 1A. Proteins (20 µg/lane) were separated on 10% SDS-PAGE gels, electrotransferred onto PVDF membranes, and probed with a monoclonal anti-LRRC8A antibody (Santa Cruz, cat. #sc-517113; 1:1,000) followed by HRP-conjugated anti-mouse secondary antibody (GE Healthcare, cat. #NA931; 1:10,000). Immunoreactivity was visualized using an ECL detection kit and imaged with a Bio-Rad ChemiDoc Imaging System. Membranes were then reprobed for β-actin as a loading control using an HRP-conjugated antibody (Millipore-Sigma, cat. #A3854; 1:50,000).

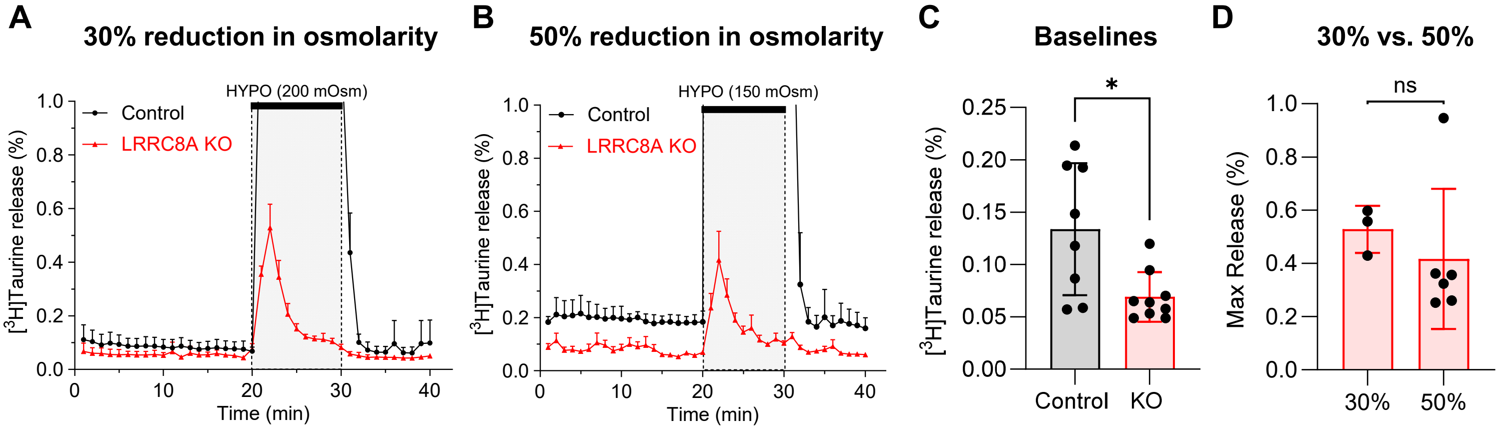


**An extended analysis of basal and swelling-activated [$^3$H]taurine release in control and LRRC8A KO astrocytes from the datasets presented in main Figure 1B and 1C**

Because small changes in unstimulated and swelling-activated [$^3$H]taurine release in LRRC8A-null cells are difficult to visualize, we present the data from main Figure 1B, C on an expanded scale. **A**, Basal [$^3$H]taurine release and the response to a 30% reduction in extracellular osmolarity from Figure 1B, shown on an expanded scale. Data are the mean values ± SD from three-four independent experiments in at least one astrocyte culture per group. **B**, Basal [$^3$H]taurine release and the response to a 50% reduction in extracellular osmolarity from Figure 1C, shown on an expanded scale. Data are the mean values ± SD from four-six independent experiments in two different astrocyte cultures per group. **C**, Statistical comparison of the baseline [$^3$H]taurine release values under isoosmotic conditions in control and LRRC8A KO astrocytes (average [$^3$H]taurine release from min 10-20). Data are the mean values ± SD from eight-nine independent experiments in at least three astrocyte cultures per group derived from panels *A* and *B*. $p<0.05$, unpaired t-test. **D**, Statistical comparison of the maximal [$^3$H]taurine release values in LRRC8A-null astrocytes in response to a 30% or 50% reduction in osmolarity. Data are the mean values ± SD from three-six independent experiments in at least one astrocyte culture per group derived from panels *A* and *B*. $p=0.512$, unpaired t-test.

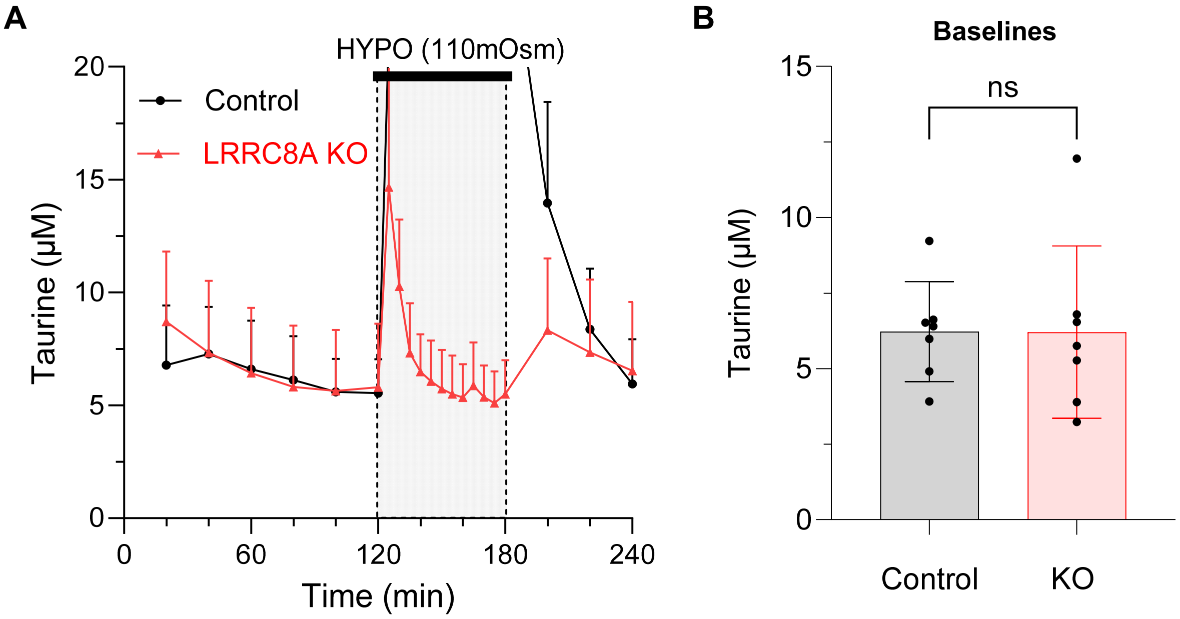


**An extended analysis of taurine levels in microdialysate samples collected in control and bLRRC8A KO brains from the datasets presented in main Figure 1D**

Because small changes in unstimulated taurine levels in LRRC8A-null brain are difficult to visualize, we present the data from main Figure 1D on an expanded scale **A,** The basal extracellular taurine levels in microdialysate samples. Data are the mean values ± SD from seven animals/genotype. **B**, Statistical comparison of the average baseline taurine levels in control and LRRC8A KO brains (averages from min 40-120). Data are the mean values ± SD from seven animals/genotype derived from panel *A***.** p=0.986, unpaired t-test.

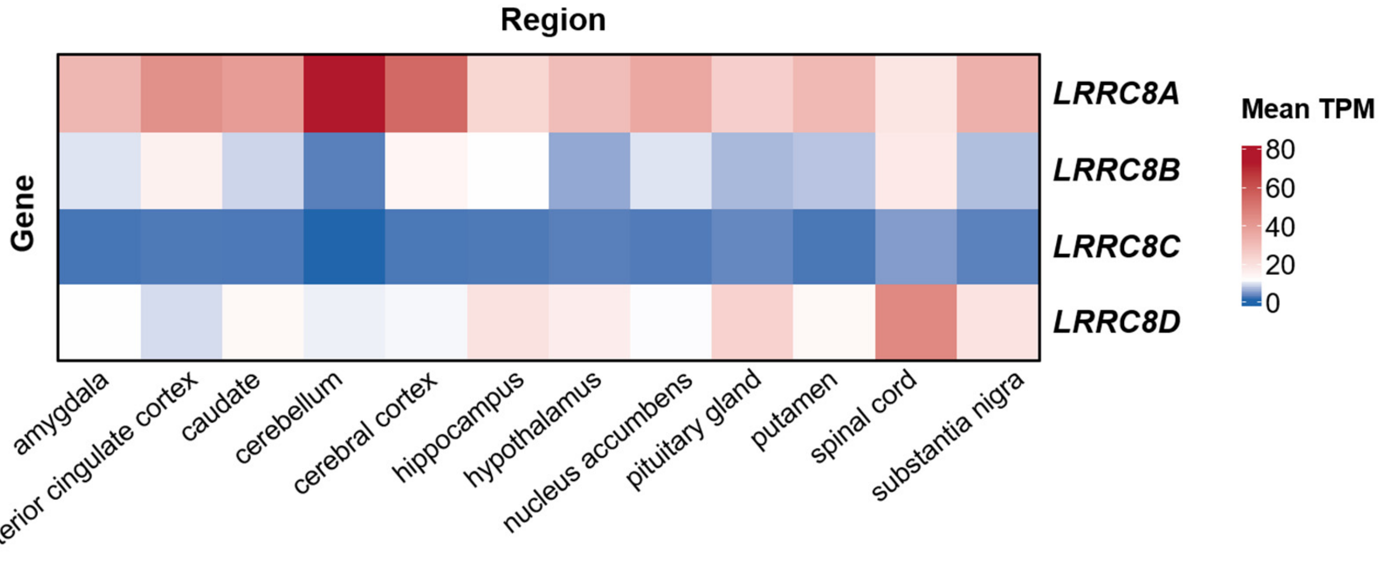


**Analysis of *LRRC8A-D* expression across 12 human brain regions**

Average TPM values for *LRRC8A-D* expression in 12 human brain regions were downloaded from the Genotype-Tissue Expression project (GTEx, https://gtexportal.org) on September 29, 2026. Selected brain regions and sample sizes were as follows: amygdala, n = 181 (52 females, 129 males); anterior cingulate cortex, n = 233 (56 females, 177 males); caudate, n = 300 (76 females, 224 males); cerebellum, n = 266 (72 females, 194 males); cortex, n = 270 (75 females, 195 males); hippocampus, n = 255 (65 females, 190 males); hypothalamus, n = 257 (64 females, 193 males); nucleus accumbens, n = 285 (73 females, 212 males); pituitary gland, n = 313 (84 females, 229 males); putamen, n = 254 (61 females, 193 males); spinal cord, n = 204 (68 females, 136 males); and substantia nigra, n = 183 (50 females, 133 males).

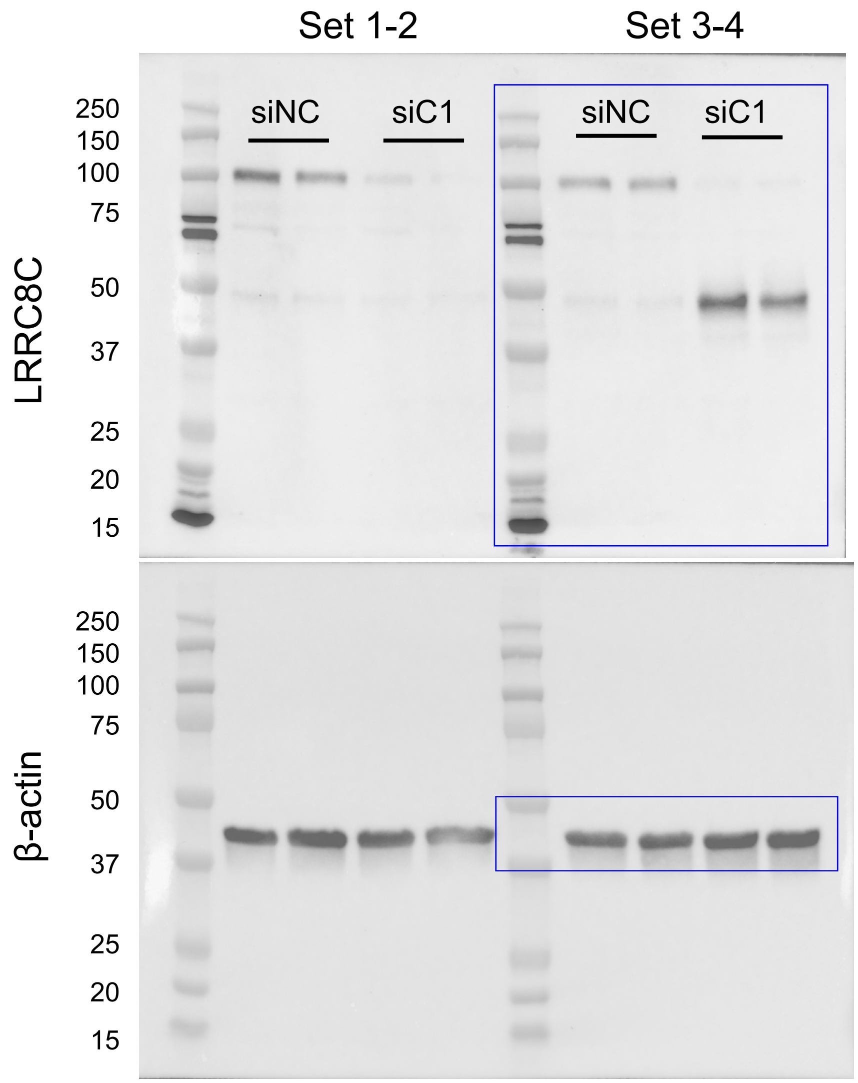


**The complete set of western blot images showing LRRC8C and β-actin immunoreactivity in four sets of protein lysates that were collected 96 hours post treatment with either negative control siRNA (siNC) or *Lrrc8c*-targeting siRNA (siC1)**

This is the full dataset of western blot images represented and quantified in the main Figure 3A-B. *Blue boxes* show the representative cropped images in the main Figure 3A. Proteins (10 µg/lane) were separated on 10% SDS-PAGE gels, electrotransferred onto PVDF membranes, and probed with a polyclonal anti-LRRC8C antibody (Proteintech, cat. #21601-1-AP; 1:2,000) followed by HRP-conjugated anti-rabbit secondary antibody (GE Healthcare, cat. #NA934; 1:10,000). Immunoreactivity was visualized using an ECL detection kit and imaged with a Bio-Rad ChemiDoc Imaging System. Membranes were then reprobed for β-actin as a loading control using an HRP-conjugated antibody (Millipore-Sigma, cat. #A3854; 1:50,000).

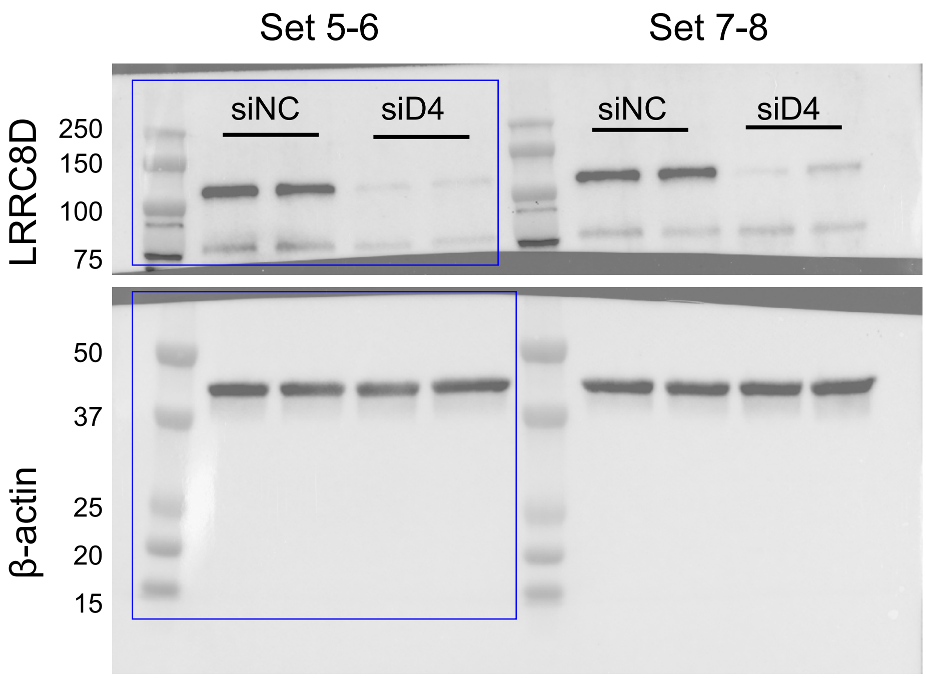


**The complete set of western blot images showing LRRC8D and β-actin immunoreactivity in four sets of protein lysates that were collected 96 hours post treatment with either negative control siRNA (siNC) or *Lrrc8d*-targeting siRNA (siD4)**

This is the full dataset of western blot images represented and quantified in the main Figure 3C-D. *Blue boxes* show the representative cropped images in the main Figure 3C. Due to a strong nonspecific signal at ~50kD, membranes were cut at ~75 kDa, with the upper portion incubated with the primary LRRC8D antibody and the lower portion incubated with an HRP-conjugated β-actin antibody. Proteins (10 µg/lane) were separated on 10% SDS-PAGE gels, electrotransferred onto PVDF membranes, and probed with a polyclonal anti-LRRC8D antibody (Proteintech, cat. #11537-1-AP; 1:2,000) followed by HRP-conjugated secondary antibody (GE Healthcare, cat. #NA934; 1:10,000). The bottom part of the membrane was probed for β-actin as a loading control using an HRP-conjugated antibody (Millipore-Sigma, cat. #A3854; 1:50,000). Immunoreactivity was visualized using an ECL detection kit and imaged with a Bio-Rad ChemiDoc Imaging System.

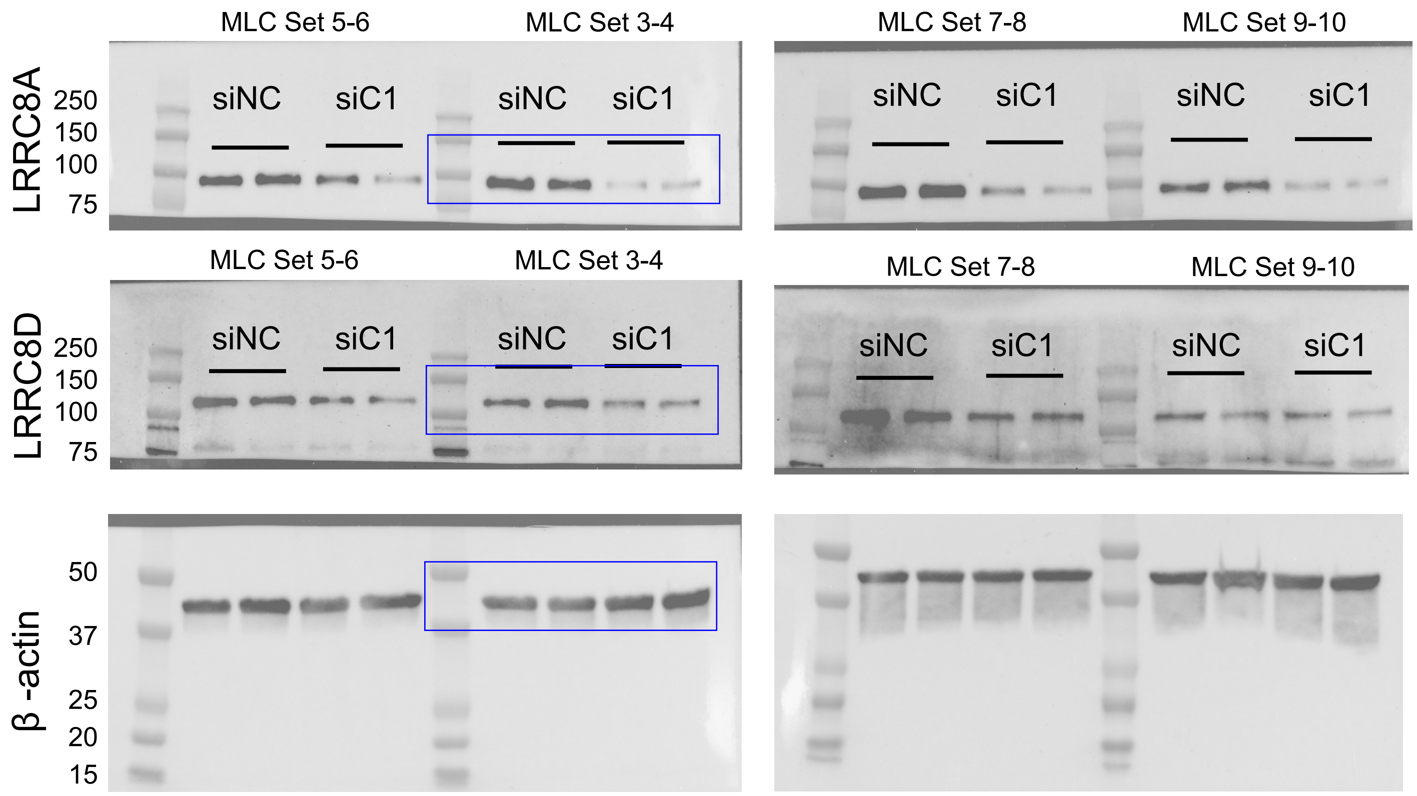


**The complete set of western blot images showing LRRC8A or LRRC8D expression in eight sets of protein lysates that were collected 96 hours post treatment with negative control siRNA (siNC) or *Lrrc8c*-targeting siRNA (siC1)**

This is the full dataset of western blot images represented and quantified in the main Figure 3E. *Blue boxes* show the representative images cropped in the main Figure 3E. The same membrane was used to probe for LRRC8A and LRRC8D given that the primary antibodies were raised in different species. Therefore, the β-actin loading control images are the same. Proteins (10 µg/lane) were separated on 10% SDS-PAGE gels, electrotransferred onto PVDF membranes, which were cut at ~75 kD. The upper part of the membrane was sequentially probed with either a monoclonal anti-LRRC8A antibody (Santa Cruz, cat. #sc-517113; 1:1,000) or polyclonal anti-LRRC8D antibody (Proteintech, cat. #11537-1-AP; 1:2,000) followed by a species matched HRP-conjugated secondary antibody anti-mouse (GE Healthcare, cat. #NA931; 1:10,000) or anti-rabbit (GE Healthcare, cat. #NA934; 1:10,000). The bottom part of the membrane was probed for β-actin as a loading control using an HRP-conjugated antibody (Millipore-Sigma, cat. #A3854; 1:50,000). Immunoreactivity was visualized using an ECL detection kit and imaged with a Bio-Rad ChemiDoc Imaging System..

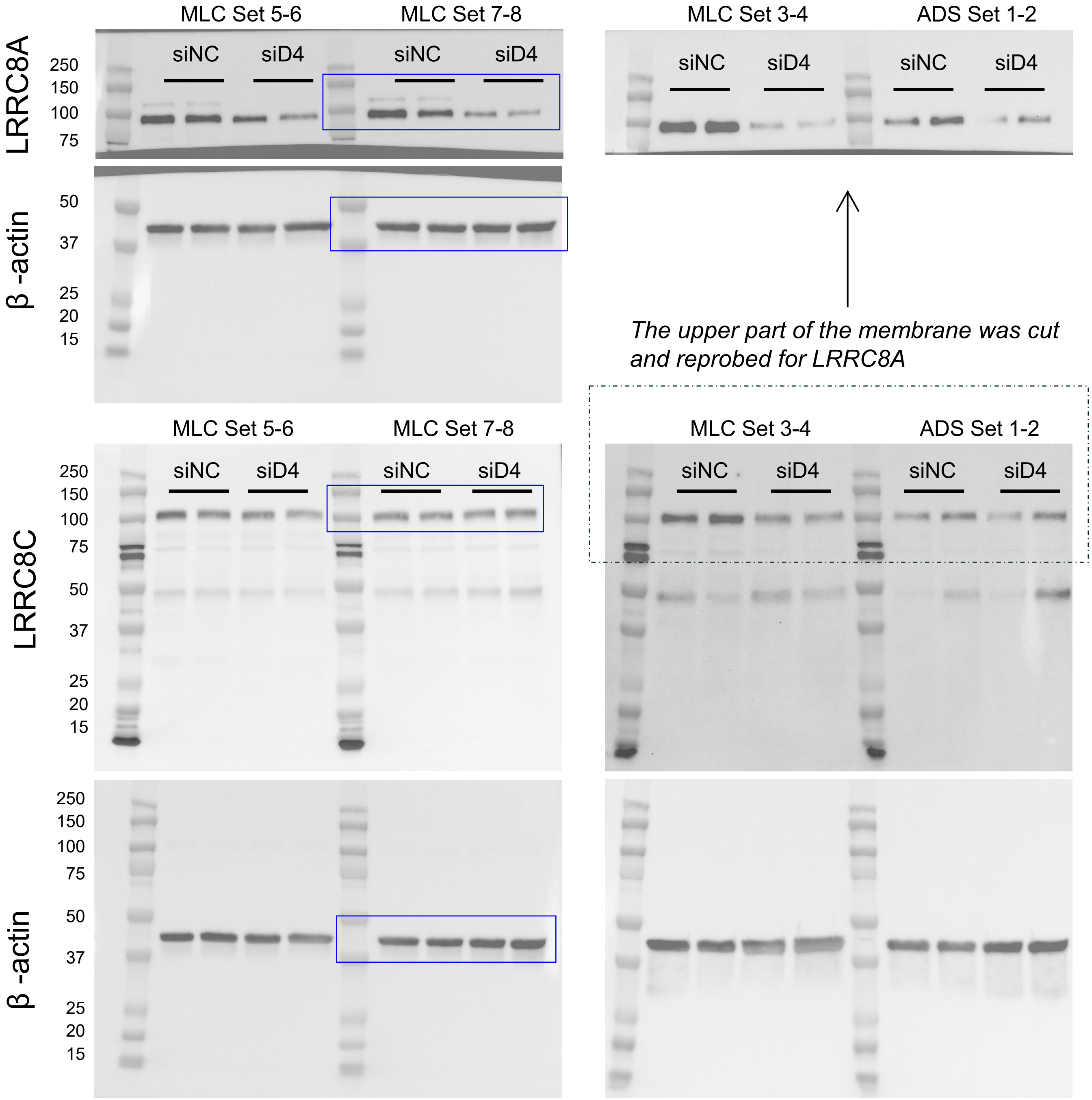


**The complete set of western blot images showing LRRC8A and LRRC8C immunoreactivity in eight sets of protein lysates that were collected 96 hours post treatment with negative control siRNA (siNC) or *Lrrc8d*-targeting siRNA (siD4)**

This is the full dataset of western blot images represented and quantified in the main Figure 3F. *Blue boxes* show the representative cropped images in the main Figure 3F. For Sets 5-8, LRRC8A was reprobed on the same membrane as in Supplemental Figure S6. Therefore, the β-actin image is the same as in Supplemental Figure S6. For images on the right, the same membrane was probed for LRRC8C and subsequently for LRRC8A. Therefore, the β-actin image is shared for both LRRC8A and LRRC8C. Proteins (10 µg/lane) were separated on 10% SDS-PAGE gels, electrotransferred onto PVDF membranes, and probed with either a monoclonal anti-LRRC8A antibody (Santa Cruz, cat. #sc-517113; 1:1,000) or a polyclonal anti-LRRC8C antibody (Proteintech, cat. #21601-1-AP; 1:2,000). This step was followed by a species-matched HRP-conjugated secondary anti-mouse (GE Healthcare, cat. #NA931; 1:10,000) or anti-rabbit antibody (GE Healthcare, cat. #NA934; 1:10,000). Immunoreactivity was visualized using an ECL detection kit and imaged with a Bio-Rad ChemiDoc Imaging System. The same membranes were probed for β-actin as a loading control using an HRP-conjugated antibody (Millipore-Sigma, cat. #A3854; 1:50,000).

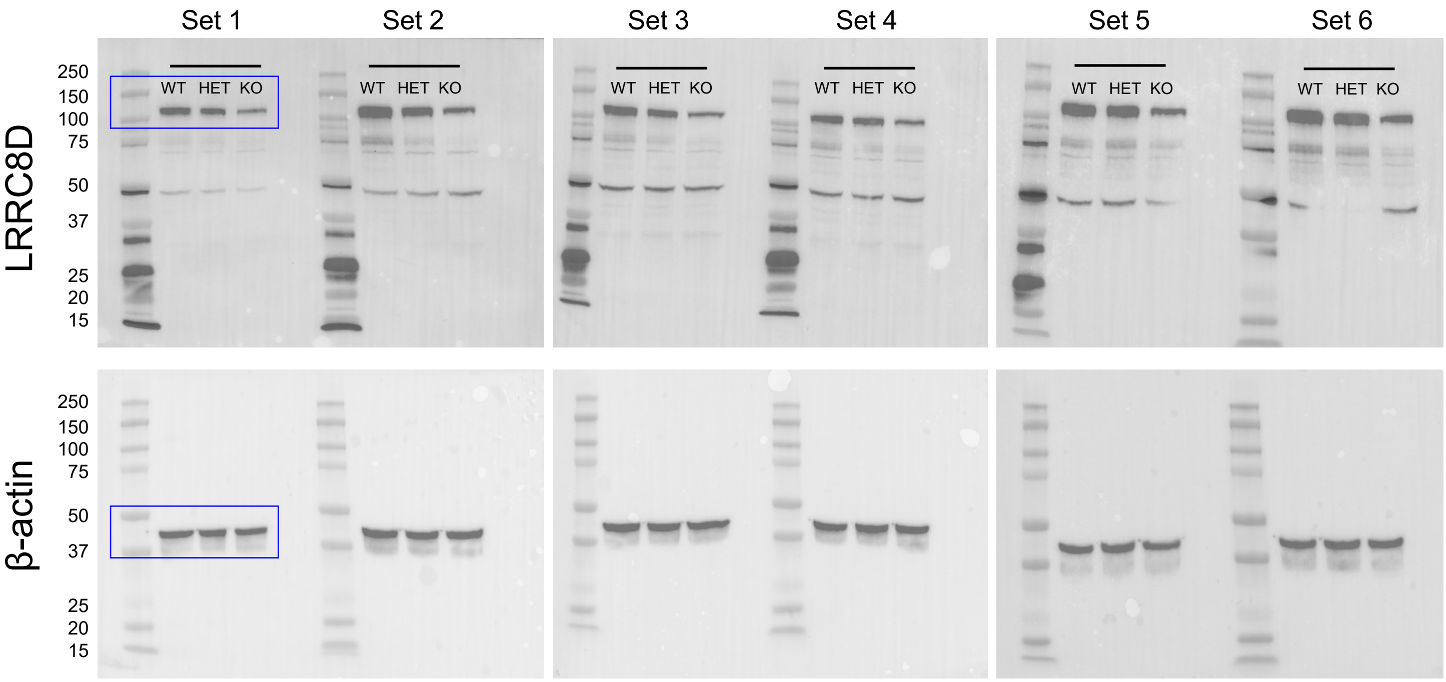


**The complete set of western blot images showing LRRC8D and β-actin immunoreactivity in six sets of protein lysates derived from LRRC8D WT, HET, and KO forebrains**

This is the full dataset of western blot images represented and quantified in the main Figure 5C. *Blue boxes* show the representative cropped images in the main Figure 5C. Proteins (20 µg/lane) were separated on 10% SDS-PAGE gels, electrotransferred onto PVDF membranes, and probed with a polyclonal anti-LRRC8D antibody (Proteintech, cat. #11537-1-AP; 1:2,000) followed by HRP-conjugated anti-rabbit secondary antibody (GE Healthcare, cat. #NA934; 1:10,000). Immunoreactivity was visualized using an ECL detection kit and imaged with a Bio-Rad ChemiDoc Imaging System. Membranes were then reprobed for β-actin as a loading control using an HRP-conjugated antibody (Millipore-Sigma, cat. #A3854; 1:50,000).

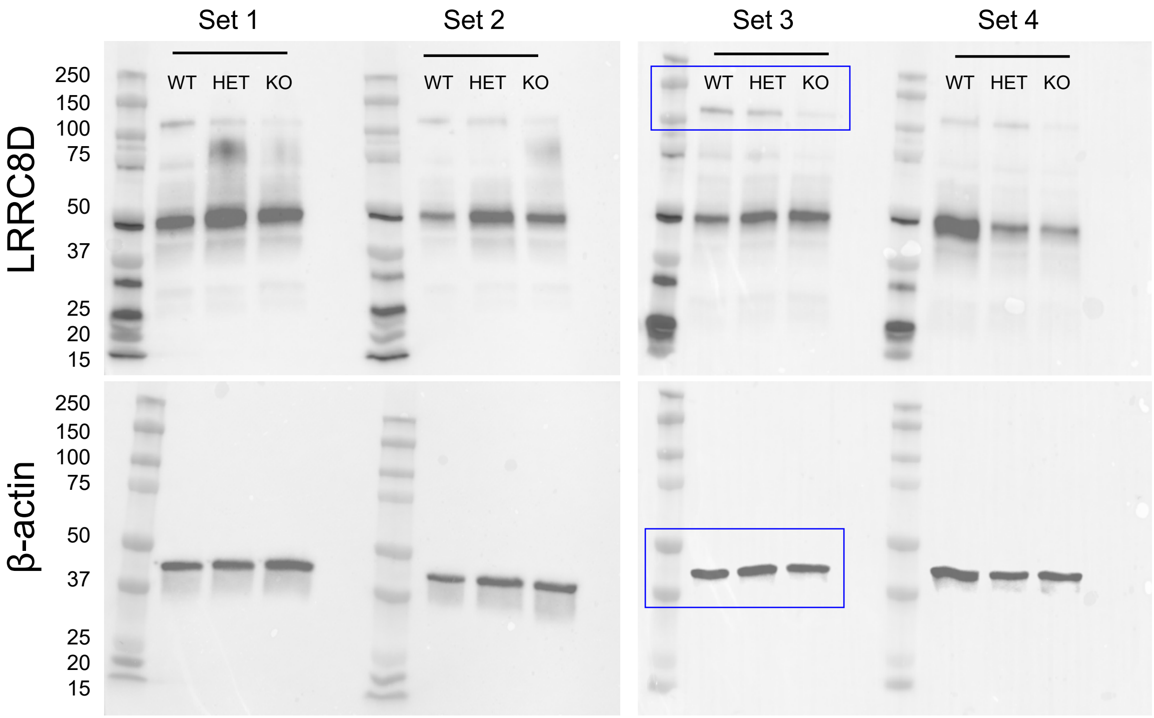


**The complete set of western blot images showing LRRC8D and β-actin immunoreactivity in four sets of protein lysates prepared from astrocyte cultures derived from LRRC8D WT, HET, and KO neonates**

This is the full dataset of western blot images represented and quantified in the main Figure 5D. *Blue boxes* show the representative cropped images in the main Figure 5D. Proteins (10 μg/lane) were separated on 10% SDS-PAGE gels, electrotransferred onto PVDF membranes, and probed with a polyclonal anti-LRRC8D antibody (Proteintech, cat. #11537-1-AP; 1:2,000) followed by HRP-conjugated anti-rabbit secondary antibody (GE Healthcare, cat. #NA934; 1:10,000). Immunoreactivity was visualized using an ECL detection kit and imaged with a Bio-Rad ChemiDoc Imaging System. Membranes were then reprobed for β-actin as a loading control using an HRP-conjugated antibody (Millipore-Sigma, cat. #A3854; 1:50,000).
